# What is in a lichen? A metagenomic approach to reconstruct the holo-genome of *Umbilicaria pustulata*

**DOI:** 10.1101/810986

**Authors:** Bastian Greshake Tzovaras, Francisca H.I.D. Segers, Anne Bicker, Francesco Dal Grande, Jürgen Otte, Seyed Yahya Anvar, Thomas Hankeln, Imke Schmitt, Ingo Ebersberger

## Abstract

Lichens are valuable models in symbiosis research and promising sources of biosynthetic genes for biotechnological applications. Most lichenized fungi grow slowly, resist aposymbiotic cultivation, and are generally poor candidates for experimentation. Obtaining contiguous, high quality genomes for such symbiotic communities is technically challenging. Here we present the first assembly of a lichen holo-genome from metagenomic whole genome shotgun data comprising both PacBio long reads and Illumina short reads. The nuclear genomes of the two primary components of the lichen symbiosis – the fungus *Umbilicaria pustulata* (*33 Mbp*) and the green alga *Trebouxia* sp. (53 Mbp) – were assembled at contiguities comparable to single-species assemblies. The analysis of the read coverage pattern revealed a relative cellular abundance of approximately 20:1 (fungus:alga). Gap-free, circular sequences for all organellar genomes were obtained. The community of lichen-associated bacteria is dominated by *Acidobacteriaceae*, and the two largest bacterial contigs belong to the genus *Acidobacterium*. Gene set analyses showed no evidence of horizontal gene transfer from algae or bacteria into the fungal genome. Our data suggest a lineage-specific loss of a putative gibberellin-20-oxidase in the fungus, a gene fusion in the fungal mitochondrion, and a relocation of an algal chloroplast gene to the algal nucleus. Major technical obstacles during reconstruction of the holo-genome were coverage differences among individual genomes surpassing three orders of magnitude. Moreover, we show that G/C-rich inverted repeats paired with non-random sequencing error in PacBio data can result in missing gene predictions. This likely poses a general problem for genome assemblies based on long reads.

## Introduction

The lichen symbiosis comprises a lichen-forming fungus (mycobiont), and a photosynthetic partner (photobiont), which is typically a green alga or a cyanobacterium. A bacterial microbiome and additional third-party fungi can also be part of the lichen consortium (Grube et al. 2015, Spribille et al. 2016). The bacterial microbiome in particular may contribute to auxin and vitamin production, nitrogen fixation, and stress protection (Erlacher, et al. 2015; Grube, et al. 2015; Sigurbjornsdottir, et al. 2016). Lichenized fungi are well known for synthesizing diverse, bioactive natural products (reviewed in (Muggia and Grube 2018)), which has recently stimulated research into biosynthetic pathways and gene clusters of these fungi (Bertrand and Sorensen 2018; Wang, et al. 2018; Calchera, et al. 2019). The estimated 17,500-20,000 species of lichens (Kirk, et al. 2008) are distributed across nearly all ecosystems (Ahmadjian 1993). Some lichens thrive as pioneering organisms in ecological niches that are otherwise adverse to eukaryotic life (Kranner, et al. 2008; Hauck, et al. 2009). The capability to inhabit such a diverse set of habitats is tightly connected with the lichen symbiosis itself. The nutritionally self-sustaining system harbors internal autotrophic photobionts, which provide carbohydrates to all other members of the association. Furthermore, some mycobiont species switch between different sets of environmentally adapted photobionts, and can thus occupy broad ecological niches (Dal Grande et al. 2018, Rolshausen et al. 2018).

There is an increasing interest in genomic resources on lichens, because lichens are valuable models in symbiosis research (Grube and Spribille 2012; Wang, et al. 2014; Grube, et al. 2015), and promising sources of biosynthetic genes for biotechnological applications (see above). Most lichenized fungi grow slowly, resist aposymbiotic cultivation, and are generally poor candidates for experimentation. Therefore, researchers increasingly use genomic data as sources of novel information on the lichen symbiosis (e.g. (Armaleo, et al. 2019)). Genome sequences of about 19 lichenized fungi and of two algal photobionts have been published to date (Table 1). Most genome sequences stem from lichens whose symbionts were grown in axenic culture. The few studies using of metagenomic data to reconstruct the fungal genomes reported highly fragmented assemblies comprising more than 1,000 scaffolds (McDonald, et al. 2013; Meiser, et al. 2017; Allen, et al. 2018). Although some assemblies range in an expected total length (McDonald, et al. 2013; Meiser, et al. 2017), and achieve comparable BUSCO (Simao, et al. 2015) scores to assemblies derived from single-species cultures (Meiser, et al. 2017), the genome sequence of *Cetradonia linearis* is to date the only publicly available genome sequence of a lichenized fungus that is based on a metagenoics approach (Allen, et al. 2018). Moreover, discontinuous assemblies are of limited use for functional genomics analyses, which rely on a comprehensive and accurate annotation of genes and even more so of gene clusters (Denton, et al. 2014; Dunne and Kelly 2017). Attempts to assemble the entire holo-genome of a lichen are missing, thus far. Also, a genome assembly strategy based on long read sequencing technology, e.g. PacBio, as well as hybrid approaches, has not yet been applied to lichens.

**Table 1.**
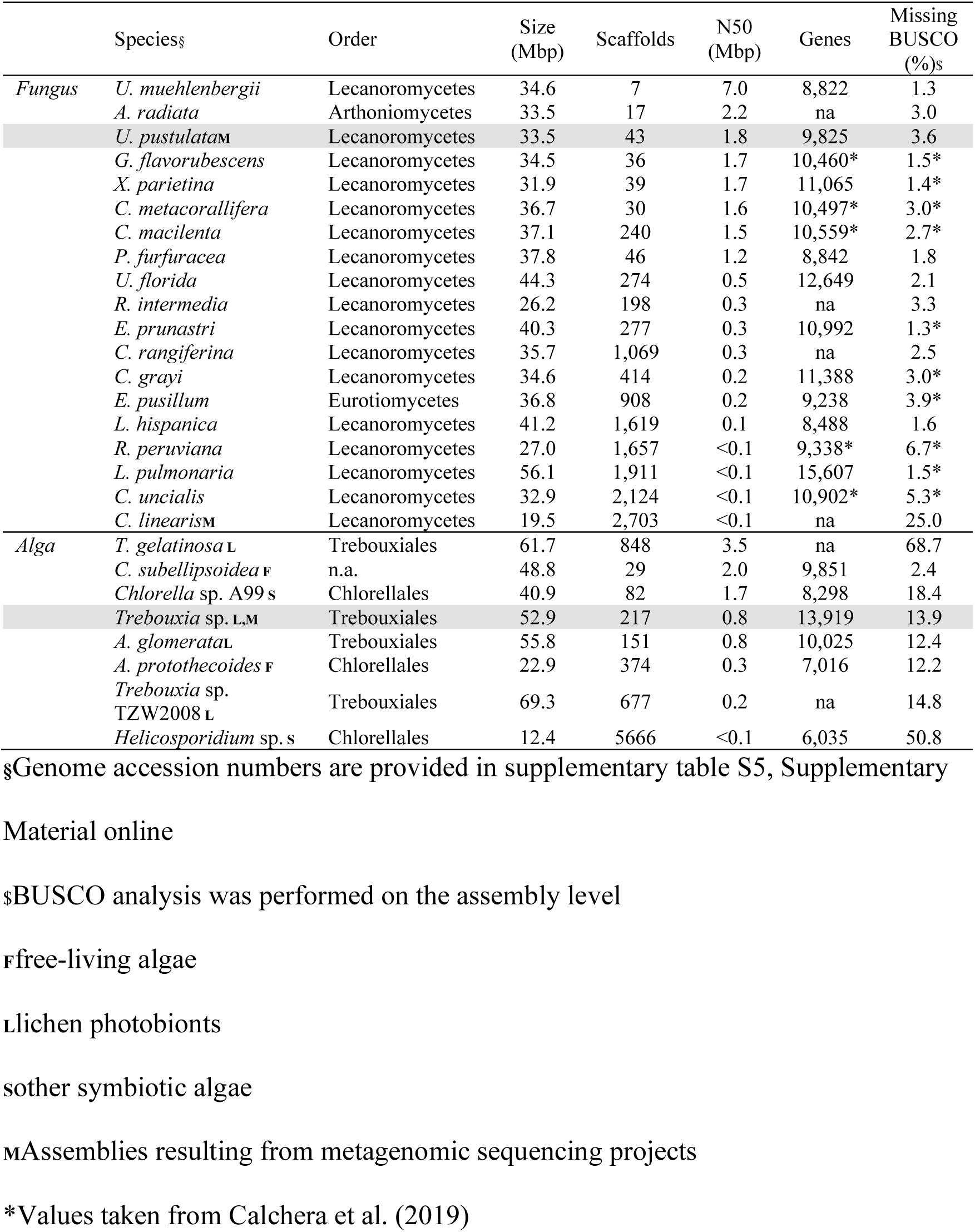
Genome assembly characteristics of a selection of lichenized fungi, and of green algae from the class *Trebouxiophyceae*. The species are sorted by descending scaffold N50. The lichen symbionts sequenced for this study are highlighted in grey.

Obtaining the complete set of genome sequences from organisms, which form obligate symbioses is challenging. Large-scale cultivation of the individual partners is often not feasible, or aposymbiotic cultivation of the symbionts is entirely impossible. This precludes efforts to obtain pure, single-species DNAs. The alternative approach, reconstructing high-quality genomes from multi-species, metagenomic samples, can be methodologically demanding (Greshake, et al. 2016). For example, genomic representation can be skewed towards one partner in the association (e.g. the host species), resulting in uneven coverage of individual genomes (Greshake et al 2016). Further methodological challenges include the risk of creating chimeric contigs, i.e. assemblies of reads from multiple genomes, or selecting the appropriate assembly software (Greshake, et al. 2016; Meiser, et al. 2017). Moreover, inaccurate post-assembly taxonomic assignment (binning) can lead to chimeric draft genome sequences, which comprise contigs from multiple species (Sangwan, et al. 2016). Thus, it is highly desirable to assess and develop methods for obtaining metagenome-assembled genomes (MAGs) of eukaryotes, and eventually achieve similar assembly qualities and reporting standards as in prokaryotes (Bowers, et al. 2017).

Here we report the reconstruction of the holo-genome for the lichen *Umbilicaria pustulata* entirely from metagenomic DNA. Details on the biology and distribution of *U. pustulata* have been published elsewhere (e,g. (Hestmark 1992; Dal Grande, et al. 2017). We inferred the genome sequences of the lichenized fungus *Umbilicaria pustulata*, its green algal symbiont *Trebouxia* sp., and its bacterial microbiome. We combined Illumina short reads from different whole genome shotgun library layouts with PacBio long reads, and integrated results from complementary assembly strategies.

Specifically, we addressed the following questions: What is the quality of fungal and algal organellar and nuclear genomes based on hybrid short and long read assemblies obtained from a metagenomic lichen sample? What are the relative genome copy numbers and the relative taxon abundances of the microorganisms involved in the lichen symbiosis? To what extent can we reconstruct the bacterial microbiome of a lichen individual? Is there evidence for horizontal gene transfer from algae or bacteria into the fungal genome? What are the methodological pitfalls associated with reconstructing the holo-genome of symbiotic communities from metagenomic reads, and with their integration into comparative genomics studies focusing on gene loss?

## Materials and Methods

### Sample collection and DNA extraction

Thalli of *U. pustulata* were collected near Olbia (Sardinia, Italy) and Orscholz (Saarland, Germany) between May 2013 and December 2014. DNA was extracted using the CTAB method (Cubero and Crespo 2002) and subsequently purified with the PowerClean DNA Clean-Up Kit (MO BIO, Carlsbad, CA, USA).

### Quantitative PCR

Quantitative PCRs (qPCR) targeted the fungal and algal single copy genes, *mcm7* (Forward - gaatgcaaggcaaacaattc, Reverse - ttgtactgttctatccgtcgg) and *g467* (COP-II coat subunit; Forward – ccttcaagctgcctatctg, Reverse - gcacctgaaggaaaagac), respectively. DNA concentrations extracted from four thalli were measured with the Qubit dsDNA High Sensitivity Kit (Life Technologies) according to the manufacturer’s instructions. For qPCR measurements, we used the *GoTaq qPCR* Master Mix (Promega) at a total volume of 10 µl. PCR (95°C for 5 min; 40 cycles of 95°C for 15 sec, 55°C for 30 sec and 60°C for 1 min) was carried out in an *ABI 7500 Fast Real Time PCR system cycler* (Applied Biosystems). Samples were measured in three technical replicates. To determine the total copy numbers, we used a standard curve approach with serial ten-fold dilutions of plasmids engineered to contain single copy PCR templates (pGEM®-T Easy Vector, Promega).

### Whole genome shotgun sequencing

We generated a whole-genome paired-end library with the Illumina TruSeq DNA Sample Prep v2 (Illumina, San Diego, CA, USA), selecting for a mean fragment length of 450 bp with the SPRIselect reagent kit (Beckman Coulter, Krefeld, Germany). A mate pair library with an insert size of 5 kb was created with the Nextera Mate Pair Sample Prep Kit (Illumina, San Diego, CA, USA). The paired-end and mate pair libraries were sequenced on an Illumina MiSeq machine. Long-read sequencing was performed on the PacBio RS II system (Pacific Biosystems of California, Menlo Park, CA, USA), using 16 SMRT cells in total.

### Read preprocessing

Low quality 3’-ends and adapter sequences were removed from the Illumina paired-end reads with Trimmomatic v0.32 (Bolger, et al. 2014) (*ILLUMINACLIP: IlluminaAdapter.fasta:2:30:10)*. Mate pairs were processed with nextclip v0.8 (Leggett, et al. 2014) to remove adapters and to bin them according to read orientation. PacBio sequence reads were error corrected with two alternative strategies. For an intrinsic error correction, we used canu v1.20 (Koren, et al. 2017). Since an intrinsic error correction requires a high long-read coverage, which might not be achieved for the less abundant genomes in the lichen holo-genome, we additionally corrected the PacBio reads using Illumina data as extrinsic information. We merged the Illumina paired-end reads with FLASH v1.2.8 (Magoc and Salzberg 2011), using standard parameters. The processed Illumina read- and mate-pair data were then assembled with MIRA v4.0, using the *genome,denovo,accurate* flags (Chevreux, et al. 1999). The resulting contigs were then used for correcting sequencing errors in the PacBio reads with ECTools (https://github.com/jgurtowski/ectools, last accessed Oct. 18 2019) requiring a minimum alignment length of 200bp with a *WIGGLE_PCT* of 0.05 and a *CONTAINED_PCT_ID* of 0.8 for the read mappings. Only PacBio reads with a length >1000 bp after correction were retained.

### De novo metagenome and metatranscriptome assembly

We employed a multi-layered strategy to target different parts of the lichen holo-genome (see supplementary text, Supplementary Material online for a detailed description of the assembly strategies and supplementary figure S1, Supplementary Material online for the workflow). In brief, we first generated an assembly of the *U. pustulata* metagenome with FALCON v0.2.1 (Chin, et al. 2016b) using the uncorrected PacBio reads. The resulting contigs were scaffolded with SSPACE-Long v.1.1 (Boetzer and Pirovano 2014). In parallel, we assembled the error-corrected PacBio reads with the Celera assembler wgs v8.3rc2 (Berlin, et al. 2015). Finally, we made a hybrid assembly with SPAdes v3.5.0 (Bankevich, et al. 2012), that made use of all Illumina reads, the ECTools error-corrected PacBio reads, and the uncorrected PacBio reads to support scaffolding. Subsequent to taxonomic assignment with MEGAN v.5.10 (Huson, et al. 2016) (see below), we binned all algal and bacterial contigs, respectively. They were then merged into single assemblies using *minimus2* (Treangen, et al. 2011) followed by a scaffolding step with SSPACE-Long with the help of the PacBio reads. For the genome of the fungus *U. pustulata*, the SPAdes contigs of at least 3 Kbp in length were used to further scaffold the FALCON assembly with SSPACE-Long. The final assemblies were polished with Pilon v1.15 (Walker, et al. 2014) using the Illumina short reads.

For the reconstruction of the organellar genomes, we used a baiting strategy. We aligned the canu-corrected PacBio reads against the organellar genomes of the lecanoromycete fungus *Cladonia grayi* (JGI Clagr3 v2.0) and the green alga *Asterochloris glomerata* (JGI Astpho2 v2.0)(Armaleo, et al. 2019) with BLAT v35 (Kent 2002), using no cut-offs. The baited reads were assembled with canu v1.20, and the resulting organellar genomes were circularized with the help of the canu-corrected PacBio reads and circlator v.1.2.0 (Hunt, et al. 2015). Assembly polishing was performed as described above.

For the reconstruction of the metatranscriptome, we assembled the RNAseq data provided in (Dal Grande, et al. 2017) with Trinity release 2013-11-10 (Haas, et al. 2013), using the *– jaccard-clip –normalize_reads* parameters.

### Taxonomic assignment

All reads and contigs were used individually as query for a DIAMOND v.0.6.12.47 search (Buchfink, et al. 2015). Contigs were searched against a custom database comprising 121 fungi, 20 plants, eight animals, 1471 bacteria and 560 viruses (supplementary table S1, Supplementary Material online), and reads were searched against the NCBI nr database. All sequences were subsequently taxonomically classified with MEGAN v5.10 (Huson, et al. 2016) requiring a minimum DIAMOND alignment score of 50. For MEGAN analyses including more than one read set, we normalized counts to the smallest read set in the analysis. Metagenomic compositions were visualized with *Krona* (Ondov, et al. 2011).

### Read mapping and coverage distribution analysis

Reads from the three WGS libraries were mapped to the assembled scaffolds with bowtie2 (Langmead and Salzberg 2012). RNAseq reads of *U. pustulata* (Dal Grande, et al. 2017) were mapped with *HISAT2* (Kim, et al. 2015), setting the maximal intron length to 3000 and keeping standard parameter values otherwise. To visualize the variation of the WGS read coverages and of the G/C content across the different genomes, we split all scaffolds into partitions of 20 Kbp in length, and subsequently clustered the individual partitions by their tetra-nucleotide frequencies. For each partition, we then plotted the mean read coverage for each WGS library, and the mean GC content with Anvi’o (Eren, et al. 2015).

### Nuclear and organellar genome annotation

Interspersed repeats were annotated with the RepeatModeler/RepeatMasker pipeline (Smit, et al. 2015). The fungal nuclear genome was annotated with funannotate (https://funannotate.readthedocs.io, last accessed Oct. 18 2019). As training data, we used the proteomes of *Xanthoria parietina* and *Cladonia grayi* (Armaleo, et al. 2019), together with *U. pustulata* transcripts. The transcripts were obtained in the following way. RNAseq data from *U. pustulata* (Dal Grande, et al. 2017) was de-novo assembled with Trinity (Haas, et al. 2013). In addition, we performed a second, reference-based assembly of the RNAseq data using Trinity’s reference-guide mode together with the fungal genome assembly. Both assemblies, together with the raw read sets, were used to identify transcripts with PASA (Haas, et al. 2008).

The nuclear genome of *Trebouxia sp*. was annotated with Maker v2.31.8 (Holt and Yandell 2011), utilizing GeneMark (Besemer and Borodovsky 2005), AUGUSTUS v3.1 (Stanke, et al. 2006), and SNAP v2006-07-28 (Korf 2004). CEGMA (Parra, et al. 2007), RNAseq data (Dal Grande, et al. 2017) and the proteome of *Asterochloris glomerata* (JGI Astpho2 v2.0) were used for model training. The organelle genomes were annotated using MFannot via the web service provided at http://megasun.bch.umontreal.ca/RNAweasel/ (last accessed Oct. 18 2019). BLAST2GO (Gotz, et al. 2008) and BlastKOala (Kanehisa, et al. 2016) were used to assign Gene Ontology terms and KEGG identifiers to the predicted genes. The graphic representation of the organellar genomes were generated with OGDraw (https://chlorobox.mpimp-golm.mpg.de/OGDraw.html, last accessed Oct. 18 2019).

### Manual curation of gene loss

To assess whether the absence of evolutionary old genes from the *U. pustulata* draft genome sequence is likely a methodological artefact or indeed indicates a gene loss, we performed a gene neighborhood analysis. In brief, we determined the ortholog to the missing LCALec gene in the close relative, *Umbilicaria hispanica* (Dal Grande, et al. 2018), and identified its flanking genes. Next, we searched for the orthologs of these flanking *U. hispanica* genes in *U. pustulata*. We decided on a methodological artefact, if any of these orthologs reside at the terminus of either a contig or a scaffold. Otherwise, we extracted the genomic regions flanking the *U. pustulata* orthologs and used it as a query of a BlastX search (Altschul, et al. 1997) against NCBI nr-prot. In addition, we used the *U. hispanica* protein as query for a tBlastN search in the *U. pustulata* genome assembly. Only when both searches provided no evidence of the missing gene, we inferred gene loss.

### Data accessibility

The raw Illumina and PacBio sequence reads have been deposited in the NCBI Sequence Read Archive (SRR8446862-SRR8446881). The *U. pustulata* and *Trebouxia sp.* assemblies have been deposited at GenBank under the accession numbers VXIT00000000 and VXIU00000000, respectively.

## Results and Discussion

### Reconstructing the holo-genome sequence of U. pustulata

Umbilicaria pustulata is a rock-dwelling lichen (Figure 1A), for which all attempts to cultivate the mycobiont in isolation have failed so far. This leaves a metagenomic approach as currently the only option to reconstruct the genome sequences of the lichen symbionts. qPCR revealed an average ratio of fungal to algal genomes in the lichen thallus of 16.2 (SD: 5.2) (supplementary table S2, Supplementary Material online). Such skewed data challenge individual assemblers to an extent that no single tool is capable to faithfully reconstruct all genomes (Bradnam, et al. 2013; Greshake, et al. 2016). We therefore, devised a sequencing and assembly scheme to reconstruct the lichen holo-genome at high contiguity (for details on the workflow see supplementary figure S1 and supplementary text, Supplementary Material online). In brief, we used both Illumina short reads and PacBio long read data, and integrated three assemblers: FALCON (Chin, et al. 2016a) for assembling uncorrected full length PacBio data; the Celera assembler (Berlin, et al. 2015) for assembling the extrinsically error corrected— and thus often fragmented—PacBio reads; and SPAdes (Bankevich, et al. 2012) for a hybrid assembly of both Illumina and PacBio reads (supplementary figure S1, Supplementary Material online). No individual method sufficed to reconstruct all genomes. A taxonomic assignment of the contigs revealed, however, that the tools complement each other in assembling different parts of the holo-genome at different contiguities (supplementary table S3, Supplementary Material online). A joint scaffolding of all fungal contigs resulted in a U. pustulata mycobiont genome sequence of 33 Mbp comprising 43 scaffolds with a scaffold N50 of 1.8 Mbp. Merging and scaffolding of the algal contigs generated 217 scaffolds with an N50 of 0.8 Mbp and a total assembly length of 53 Mbp. The assembly lengths for both the fungal and the algal genomes fall well in the diversity of other lichenized fungi and members of the Trebouxiophyceae, respectively (Table 1). Merging and scaffolding the bacterial fraction of the three assemblies resulted in 499 contigs amounting up to 35 Mbp. Two bacterial contigs with lengths of 3.6 and 3.4 Mbp, respectively, represent major parts of two genomes from the genus Acidobacterium. We refer to them as Acidobacterium C16 and Acidobacterium C35, respectively.

**Fig. 1.**
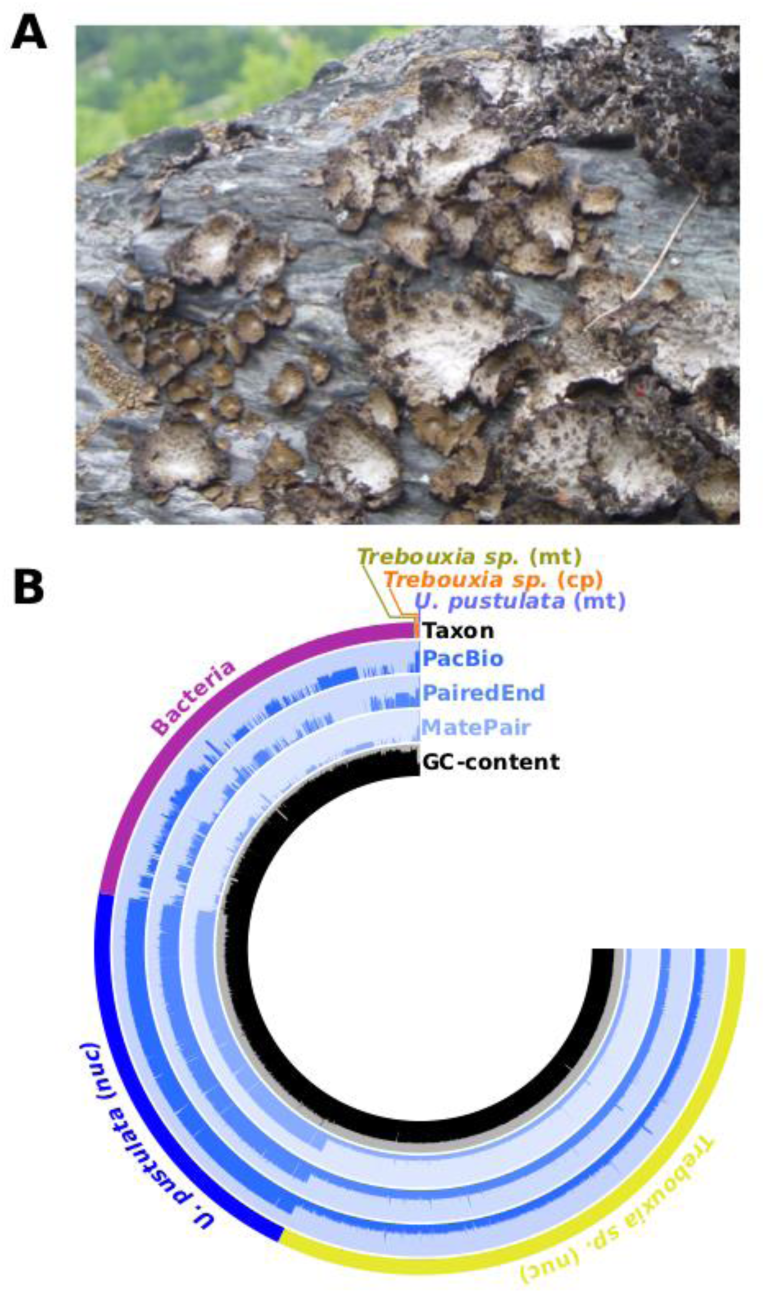
**A** The lichen *Umbilicaria pustulata*. **B** Read coverages and G/C content distribution across the genomes in the lichen holo-genome and the three whole genome shotgun libraries. Bar height indicates mean read coverage or mean G/C content in a 20 kb window of the corresponding assembly on a log scale. The fungal mitochondrial genome of the fungus (*U. pustulata* (mt)) is, with a mean read coverage (PacBio) of 3,713, the most abundant component of the holo-genome.

No scaffold in the final assembly represented the full-length genomes of the fungal and algal mitochondria, or of the algal chloroplast. We therefore used the organellar genome sequences of Cladonia grayi and of Asterochloris glomerata as baits to identify PacBio reads originating from the organellar genomes. The baited reads were assembled individually for each genome, resulting in a circular, gap-free sequence for each of the three organelles (supplementary figures S2-S4, Supplementary Material online). The fungal mitochondrial genome (mt genome) comprises 95.4 kb. It ranks third in length among 22 mt genomes from lecanoromycete lichens (Pogoda, et al. 2018), superseded only by Leptogium hirsutum (120 kbp) and Parmotrema stuppeum (109 kbp). The algal mitochondrion and chloroplast have lengths of 99.9 kbp and 272.0 kbp, respectively. They are larger than the organellar genomes in other Trebouxiophyceae, both symbiotic and free living (Fan, et al. 2017).

### Taxon abundance in the lichen holo-genome

The metagenomic reconstruction of the lichen holo-genome allows, for the first time, to infer average genome copy numbers in a lichen thallus from the read coverage distribution. Figure 1B and supplementary table S4, Supplementary Material online, reveal that the coverage variation between the various genomes extends over three orders of magnitude. The coverage for the fungal nuclear genome assembly, and thus the genomic copy number, is on average about 20 times (SD: 5.9) higher than that of the algal nuclear genome assembly, which corroborates the findings from the qPCR analysis. Since both symbionts are haploid, this translates into an average abundance of 20 (SD: 5.9) fungal cells per algal cell. Each fungal cell harbours 15 (SD: 4.2) copies of the mitochondrial genome. In each *Trebouxia sp.* cell, there are 17 (SD: 6.2) copies of the mitochondrial genome. *Trebouxia sp*. possesses only a single chloroplast. Thus, similar to many other green microalgae (Gallaher, et al. 2018), the *Trebouxia sp.* chloroplast genome is polyploid and contains, on average, 16 (SD: 5.2) copies. To our knowledge, this is the first report of ploidy level for the chloroplast in a lichenized green alga. The two *Acidobacterium spp.* are each represented with about one cell per algal cell.

### Characterization of the bacterial community

To further analyze the taxonomic composition of the bacteria that are associated with *U. pustulata*, we performed a taxonomic assignment at the read level (Figure 2 and supplementary figure S5, Supplementary Material online). *Acidobacteriaceae*, *Alphaproteobacteria*, and *Actinobacteria* are the three main bacterial phyla, similar to what has been observed for Antarctic lichens (Park, et al. 2016). In general, the taxonomic composition resembles closely typical rock-inhabiting bacterial communities (Choe, et al. 2018). Yet, other studies suggested that *Alphaproteobacteria* and not *Acidobacteria* dominate lichen microbiomes (e.g. (Grube, et al. 2009; Bates, et al. 2011; Aschenbrenner, et al. 2014)), with abundances of up to 32% for the *Rhizobiales* in the lichen *Lobaria pulmonaria* (Erlacher, et al. 2015). This may indicate that microbiome compositions can vary considerably between lichen species. However, differences in the methodology for assessing taxon frequencies can also result in substantially deviating results (Nayfach and Pollard 2016). The microbiome analyses by Erlacher, et al. (2015) were performed at the level of assembled contigs. While this eases the taxonomic assignment, due to the use of longer sequences (Vollmers, et al. 2017), it is bound to result in distorted abundance estimates. The high read coverage for abundant taxa in a microbiome generally results in more contiguous assemblies comprising only few contigs. In a typical MEGAN analysis, taxon abundance is assessed by the number of sequences that are assigned to that taxon. As a consequence, common taxa with contiguous genome assemblies will receive low counts, and their abundance will be underestimated. Rare taxa, in turn, whose lower read coverage results in more fragmented genome reconstructions with many short contigs will receive high counts. Their abundance will be overestimated (supplementary figure S6, Supplementary Material online). We demonstrate the effect of the chosen methodology on the reconstruction of the *U. pustulata* microbiome (supplementary figure S7, Supplementary Material online). Applying the method of Erlacher, et al. (2015) increased the estimated abundance of the *Rhizobiales* to 11%, and decreased that of the *Acidobacteriaceae* to 18%. The dominance of the *Acidobacteriaceae* was restored when pursuing a hybrid approach, in which the taxonomic assignment was done at the contig level and the abundance estimates were based on the reads mapping to the contigs (supplementary figure S7B, Supplementary Material online). We conclude that the methodological impact on the taxon abundance estimates is substantial, and needs to be taken into account when comparing microbiome community composition in different studies.

**Fig. 2.**
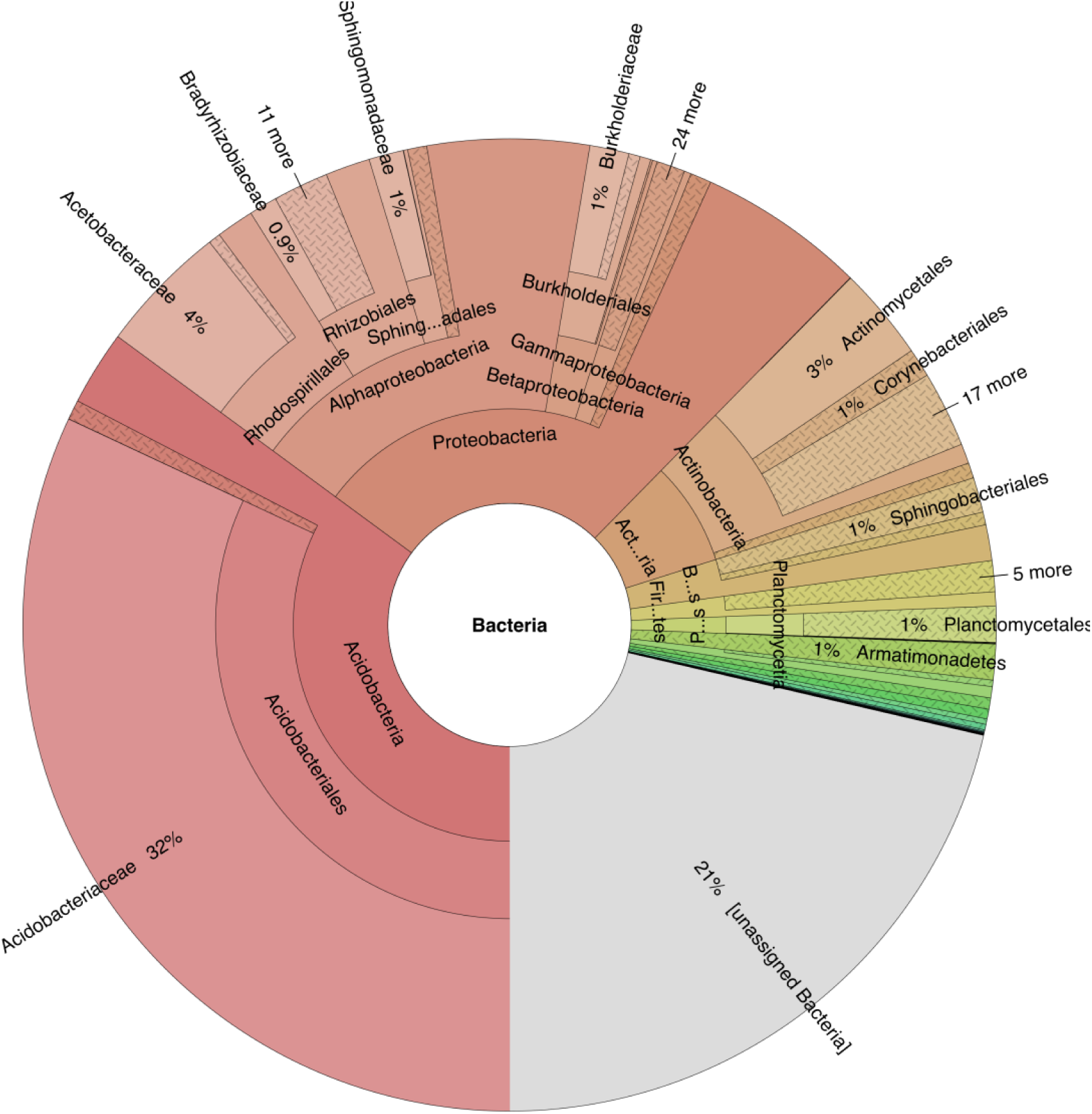
Composition of the bacterial fraction represented in the *U. pustulata* metagenomic reads. Reads from the two Illumina whole genome shotgun libraries and the PacBio reads were pooled and taxonomically assigned with MEGAN (Huson, et al. 2016). *Acidobacteria*, uniting 35% of the read counts, *Proteobacteria* (27%), and *Actinobacteria* (8%) are the three most abundant phyla. Notably, a single family, the *Acidobacteriaceae* (32%), dominates the microbiome. Its most abundant genera are *Granulicella*, *Terriglobus*, and *Acidobacterium* to which the two largest bacterial contigs belong to. Among the *Proteobacteria*, *Rhodospirillales* (6%), and therein the *Acetobactereaceae* (4%) take the largest share, followed by the *Rhizobiales* (4%). Within the *Actinobacteria*, *Actinomycetales* are the dominant family (3%). See supplementary figure 5, Supplementary Material online, for a species-level resolution of the microbiome.

### Annotation of the nuclear genomes

The nuclear genome of *U. pustulata* (mycobiont) has an average G/C content of 51.7%, and interspersed repeats account for 25.5% of the sequence. We identified 9,825 protein-coding genes (Table 1), with on average 3.3 exons, and a mean transcript length of 1,406 bp. A BUSCO analysis (Simao, et al. 2015) revealed that 94.4% of 1,315 the genes in the ‘Ascomycota’ dataset are represented over their full length in the genome sequence. This value is in the same range of what is typically achieved for fungal genomes reconstructed from axenic cultures (Table 1, supplementary table S5, Supplementary Material online).

The genome of *Trebouxia* sp. has an average G/C content of 50.0%, and interspersed repeats account for only 4.9% of the sequence. We predicted 13,919 genes with on average 6.7 exons per gene, and a mean transcript length of 1,221 bp. With 13.9%, the fraction of genes from the ‘Chlorophyta’ BUSCO (2,168 genes) that were not found in the genome sequence is considerably high. However, similar results are obtained when analyzing other representatives of the *Trebouxiophyceae* with both free living and symbiotic lifestyles (Table 1). A notable exception, with only 2.4 % missing BUSCOs, is *Coccomyxa subellipsoidea*. This is, however, not surprising since this species was used for the initial compilation of the ‘Chlorophyta’ BUSCO set. We have shown previously that even highly fragmented genome assemblies can recover most of the BUSCO genes (Greshake, et al. 2016). Thus, our results indicate that the plasticity of the algal gene set might be higher than hitherto acknowledged.

### No evidence for horizontal gene transfer in U. pustulata

The lichen symbiosis, an evolutionarily old, obligate, and stable association of individuals from different species, should provide an optimal basis for the mutual exchange of genetic material. We therefore screened the fungal genome assembly for indications of horizontal acquisitions of either algal or bacterial genes. Ten fungal genes were classified as of algal, and further 12 as of bacterial origin. All genes are located amidst fungal genes in the genome assembly. However, a subsequent case-by-case curation of these 22 genes revealed that the taxonomic assignments by MEGAN are, in all instances, borderline cases (supplementary table S6, Supplementary Material online). The sequence similarity of the corresponding genes to an algal or bacterial protein, which served as basis for the classification, was low, and only slightly higher than the similarity to the closest fungal gene. Only a slight shift in the parameterization of MEGAN’s taxonomic classification algorithm left these genes essentially taxonomically unassigned. Thus, the true evolutionary origin remains unknown for all 22 genes. Individual examples of genetic exchange between lichenized fungi and their algal partners have been reported before (e.g. (Wang, et al. 2014; Beck, et al. 2015)). Here, we find no convincing evidence for the horizontal acquisition of either algal or bacterial genes by *U. pustulata*.

### Lineage specific absence of evolutionarily old genes in U. pustulata

We subsequently increased the resolution of the gene set analysis to search for 9,081 genes that were present in the last common ancestor of the *Lecanoromycetes* (LCALec; see supplementary text, Supplementary Material online). For 142 LCALec genes were missing an ortholog only in the *U. pustulata* gene set, suggesting, on the first sight, an exclusive loss on the *U. pustulata* lineage. On closer scrutiny, however, all but 33 of these genes had been either missed during genome annotation, or reside in assembly gaps since an ortholog could be detected in the transcript data. A corresponding analysis in four other genomes of lichenized fungi obtained similar results (supplementary text, supplementary table S7 and supplementary figure S8, Supplementary Material online).

Taking the absence of genes in annotated gene sets at face value can, therefore, lead to wrong evolutionary inferences (Deutekom, et al. 2019). However, for 33 LCALec genes we could find, to this point, no indication of an experimental artefact, and they appear genuinely absent from the *U. pustulata* genome (supplementary table S8, Supplementary Material online). Four of these genes are represented by an ortholog in the closely related *Umbilicaria hispanica* (Dal Grande, et al. 2018), dating their putative loss to after the split of the two *Umbilicaria* species. In three cases, a subsequent manual curation corroborated the gene loss assumption. The three genes encode an oxidoreductase with a significant sequence similarity to gibberellin-20-oxidases, a putative methyl-transferase, and a protein with unknown function. For one gene, DHFR, however, our curation revealed an error source in the gene identification, which is typically neglected. DHFR encodes a protein, which is involved in the basal nucleotide metabolism. This gene is almost ubiquitously present throughout fungi and animals. Its absence in *U. pustulata* therefore would imply far-reaching changes in metabolism (Huang, et al. 1992). Our manual curation could exclude assembly errors and genomic rearrangements as likely explanations for the absence of DHFR (Figure 1C). A tBlastN search with the *Saccharomyces cerevisiae* DHFR (Uniprot-ID: P07807) as query obtained a partial hit in this region, which indicated that the ORF of DHFR is disrupted by several frameshift mutations. Because this region is covered by about 200 PacBio reads, sequencing errors appeared unlikely suggesting a recent pseudogenization of DHFR in the lineage leading to *U. pustulata*. However we noted a very low Illumina read coverage at the DHFR locus (Figure 3). This coverage drop coincides with an extraordinary high G/C content of up to 78% paired with the presence of extended stretches of self-complementarity (Figure 4). In combination, this can lead to the formation of stable stem-loops that can interfere with both DNA amplification and sequencing (Benjamini and Speed 2012; Ross, et al. 2013; Schirmer, et al. 2016). We suspected that the low Illumina read coverage rendered assembly polishing with Pilon less effective. Indeed, a manual inspection exploiting the few Illumina reads that map to the DHFR locus, identified six of eight frameshift mutations as recurrent sequencing errors in the underlying PacBio reads (supplementary figures S9-S14, Supplementary Material online). The remaining two frameshifts towards the 3’-end of the ORF, which are not covered by any Illumina reads, coincide with runs of Gs. Thus, they are very likely to be also sequencing errors (supplementary figures S15-S16, Supplementary Material online). Correcting all frameshifts resulted in an uninterrupted ORF (supplementary figure S17, Supplementary Material online) encoding a full-length DHFR.

**Fig. 3.**
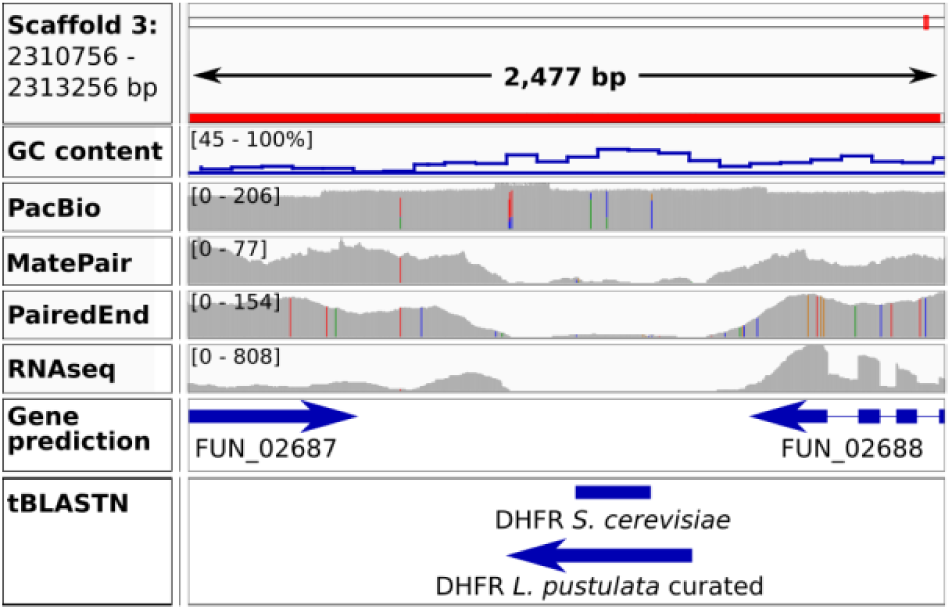
Read coverage distribution in the DHFR locus. Coverage pattern at the DHFR locus (Scaffold 3: 2,310,756-2.313,256). While the read coverage is consistently high for PacBio (∼200x), there is a marked decrease for the two Illumina whole genome shotgun libraries towards the center of this region. This decrease coincides with a marked increase of the GC content up to 79%. A tBlastN search using the Dihydrofolate Synthase of *S. cerevisiae* (Uniprot-ID: P07807) obtains a partial hit in the central part of region. Eight frameshift mutations in the CDS of DHFR were manually corrected (supplementary figures S6-S13, Supplementary Material online) resulting in a curated putative protein of 210 aa in length.

**Fig. 4.**
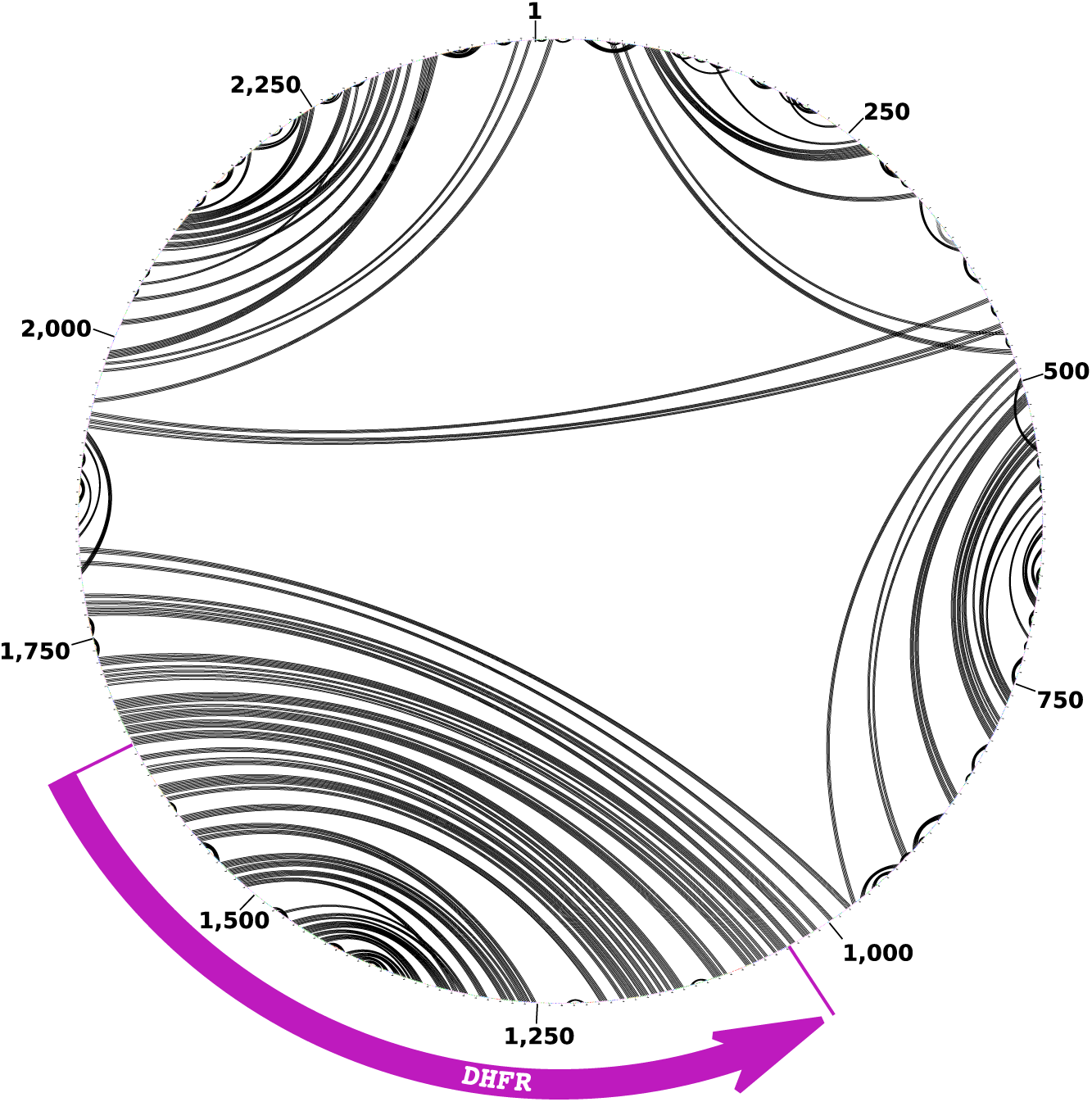
Inverted repeats in the DHFR locus. We assessed the potential of the DHFR locus to form secondary structures that may interfere with the Illumina sequencing technology. The plot shows self-complementarity predicted by ProbKnot (Bellaousov and Mathews 2010) as black arcs. The pattern reveals that the DHFR gene in *U. pustulata* is embedded in an inverted repeat spanning approximately 800 bp.

To assess the extent to which G/C-rich inverted repeats may interfere in general with the correct identification of genes, we annotated inverted repeats throughout the genome draft sequence of *U. pustulata* with the Inverted Repeat Finder (Warburton, et al. 2004). This revealed 1,464 IR, with a median length of 819.5 bp. The G/C content of these repeats follows a bimodal distribution peaking at 51% and 75%. While the number of inverted repeats falls within the values obtained for other genomes of lichenized fungi, IRs with a G/C content of over 70% are unique to *U. pustulata* (Figure 5). Whether this is due to the fact that only *U. pustulata* was sequenced with a long read technology that is less sensitive to G/C rich inverted repeats, or whether the other genomes are devoid of such repeats remains to be determined. Overlaying the IR regions with the Illumina and the PacBio read coverage information reveals 467 IR with a mean G/C content of 67.8% for which the Illumina read coverage drops to <10x while the PacBio coverage remains uniformly high. Any gene residing in such a region has a considerable chance to be either incorrectly predicted or overlooked due to remaining sequencing errors in the genome draft sequence.

**Fig. 5.**
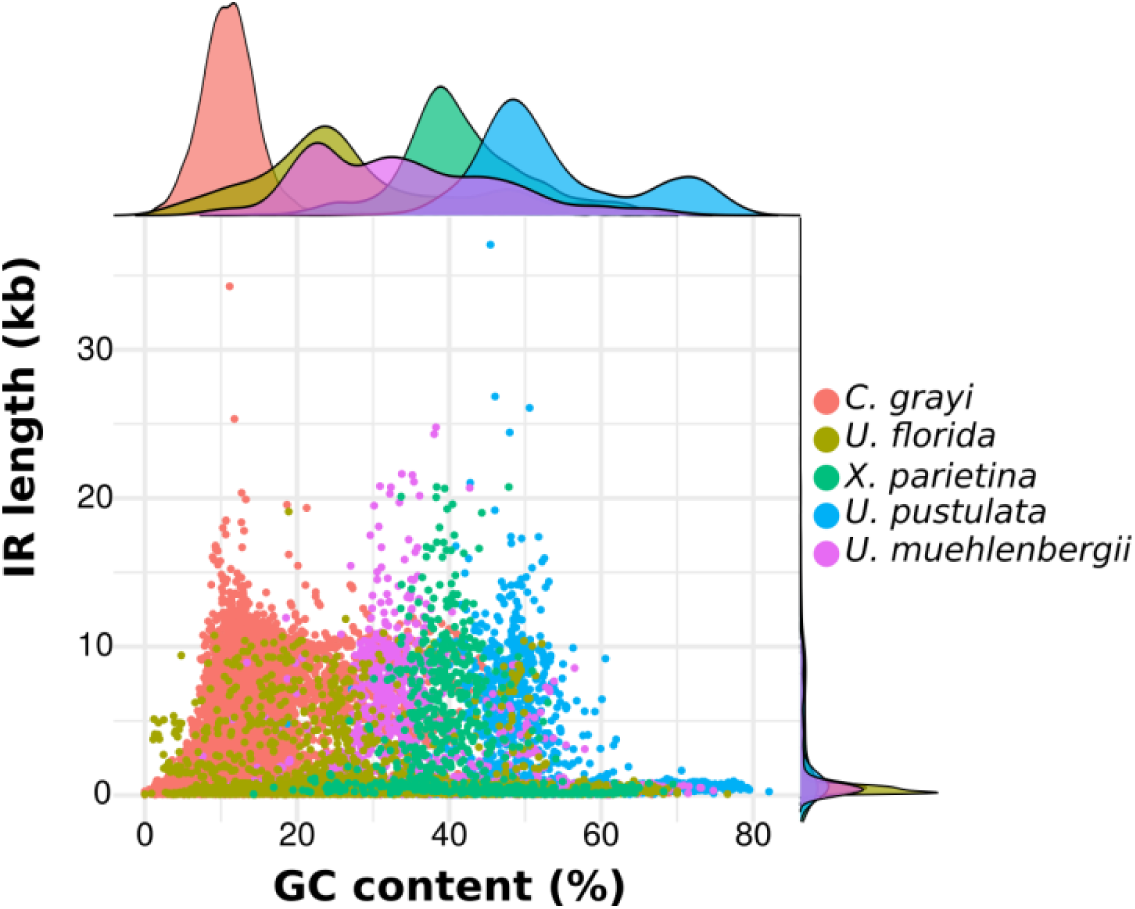
The distribution of inverted repeats in the draft genome sequences of five lichenized fungi. Inverted repeats with a GC content above 70% are observed only in *U. pustulata*.

### Organellar genome annotation

Annotation of the *L. pustulata* mitochondrial genome resulted in 15 protein-coding genes, a small subunit rRNA gene, 33 additional ORFs, and 31 tRNA genes encoding 24 distinct tRNAs (supplementary figure 2, Supplementary Material online). All 15 fungal core protein coding genes (Pogoda, et al. 2018) are represented, among them atp9, which was found to be frequently missing in the mt genomes of lichenized fungi (supplementary table S9, Supplementary Material online). While this suggests, on the first sight, a considerably standard layout of the mt genome, a closer look at the annotated genes revealed a number of interesting findings. Most notably, *cox2*, the gene encoding the cytochrome c oxidase subunit II is fused head-to-tail to *cob*, which encodes cytochrome b, into one transcription unit (supplementary figure 18, Supplementary Material online). The corresponding Trinity transcript contains an uninterrupted reading frame, suggesting that it is translated into a single fusion protein. To the best of our knowledge, such a fusion as never been reported before, although at least the lecanoromycete *Usnea ceratina* contains a similar fusion (NCBI Gene ID: 34569213). Future studies will have to reveal when during evolution this gene fusion emerged, and at what stage during gene expression—and via what mechanism—the two proteins are separated. Moreover, we noted that nad6, the gene encoding the NADH dehydrogenase subunit 6, is disrupted by the integration of a 2.4 kb long segment, most likely a mobile Group II intron (Lambowitz and Belfort 1993) (supplementary figure S19, Supplementary Material online). Eventually, three protein-coding genes do not possess a recognizable stop codon (supplementary table S9). One example is the gene encoding the NADH dehydrogenase subunit 3 (nad3). The predicted ORF is covered by three distinct transcripts, indicating that it is not a single transcription unit (supplementary figure S20, Supplementary Material online). A search against the MitoFun database (http://mitofun.biol.uoa.gr, last accessed Oct. 18 2019) reveals that the CDS encoding nad3 spans approximately the first 396 bp of this ORF. In this region, no canonical stop codon is detected, and the agreement between the transcript sequence and the genomic sequence suggests that no stop codon is generated post-transcriptionally via RNA editing. BlastP and BlastN searches (Altschul, et al. 1997) against the NCBI databases *nr-prot* and *nr*, respectively, revealed no significant hits for the parts of the ORF downstream of nad3. The absence of recognizable stop codons in the gene encoding nad3 can be found in the mt genome annotations of other *Lecanoromycetes*, e.g. in *Usnea mutabilis* (NCBI GeneID: 38289161) and *Parmotrema ultralucens* (NCBI GeneID: 38466336). It remains unclear how lichenized fungi achieve an accurate termination of the translation for such genes. Of the remaining 36 ORFs annotated in the *U. pustulata* mt genome, 9 encode homing endonucleases that have been proposed to act as selfish genetic elements driving changes in both mt genome size and gene order (Aguileta, et al. 2014; Kanzi, et al. 2016).

The annotation of the *Trebouxia sp.* mitochondrial genome revealed 32 protein coding genes, 20 additional ORFs, and 26 tRNAs, which agrees with previous findings in the *Trebouxiophyceae* (Fan, et al. 2017). In the chloroplast genome, we could annotate 78 protein coding genes, three ribosomal RNAs, 52 additional ORFs, and 31 tRNA. The set of annotated genes comprises all green algal core genes, and additionally 15 out of 16 common algal chloroplast genes showing sporadic lineage specific gene loss (Turmel, et al. 2015). Interestingly, the missing ribosomal protein, rps4, is encoded on scaffold 44 of the algal nuclear genome assembly. Here, it is flanked by two genes, whose counterparts in other green algae are located in the nucleus (supplementary figure S21, Supplementary Material online), and the read coverage pattern provides no hint for any assembly error. This indicates a recent relocation of rps4 from the chloroplast to the nucleus in *Trebouxia sp*..

### Conclusion

Here, we have shown that the reconstruction of the holo-genome for an obligate symbiotic community purely from metagenomic sequence reads at contiguities comparable to assemblies for single-species samples is feasible. The greatly varying coverage ratios for the individual genomes, spanning three orders of magnitude, emerged as the most challenging task. Key to success was the combination of short Illumina and long PacBio reads with a comprehensive assembly scheme. In particular, we had to (i) target different components of the holo-genome with different assembly methodologies, (ii) include taxonomic assignments on the contig level, (iii) perform a merging of contigs from different assembly approaches that were assigned to the same taxonomic group, and (iv) perform a final scaffolding step. Numerous benchmark studies have indicated that there is no general gold standard for a genome assembly procedure (Dominguez Del Angel, et al. 2018). Thus, our workflow should be considered a template that can be adapted to the needs of the precise symbiotic community under study. The initial analysis of the *U. pustulata* holo-genome already revealed a number of genetic changes both in the nuclear and in the organellar genomes whose functional relevance for this obligate lichen symbiosis will be interesting to determine. However, we encountered also a number of pitfalls that, if remain unnoticed, lead to wrong conclusions. One of the main advantages of metagenomic approaches is that holo-genome reconstruction, relative genomic copy number assessment, taxonomic classification and relative taxon abundance estimation will be performed on the same data. It is tempting to use the assembled contigs for the taxonomic assignments, because longer sequences will allow a classification with greater confidence. If the aim is, however, to assess the abundance of individual taxa in microbial community, the analysis has to take the read data into account. Either by performing the taxonomic assignment at the read level—bearing the risk that a fraction of reads will remain unclassified—or by taking the read coverage of the taxonomically assigned contigs into account, which will miss rare taxa covered by only few reads. From an evolutionary perspective, the availability of genome sequences for an obligate symbiotic community is the relevant starting point for determining the genetic changes underlying the dependency of the symbionts. A comprehensive gene annotation is essential for such analyses, which have a strong focus on detecting loss of individual genes. BUSCO analyses provide an initial indication for the completeness of gene annotations. However, BUSCO sets are, compared to the gene set of a species, typically small, and they are often not designed for the phylogenetic clade under study. On the example of the green algae, we showed that the latter aspect makes it difficult to differentiate between the absence of BUSCO genes due to an incomplete gene set reconstruction, or due to lineage specific losses of BUSCO genes in higher than expected numbers. The use of tailored core gene sets for the clade of interest, paired with targeted ortholog searches both in the annotated gene set and in the assembled transcriptome data is an alternative that substantially increases resolution. Genes that then remain undetected are good candidates for a lineage-specific loss with all its consequences for the symbionts’ metabolism. Still, this does not exclude an artefact. It was only the suspicious deviation in coverage between the PacBio reads and the Illumina reads, which eventually revealed that the gene encoding the dihydrofolate reductase (DHFR) was not lost in *U. pustulata*. Ultima ratio remains, therefore, expert candidate curation considering all evidences that can hint towards an artefact mimicking gene loss.

## Acknowledgements

The authors thank the LOEWE-Centre TBG funded by the Hessen State Ministry of Higher Education, Research and the Arts (HMWK), Anjuli Calchera (Frankfurt) for technical assistance, and Pavel Škaloud (Dpt. Botany, Charles University, Prague) for useful discussion on *Trebouxia* organellar genomics and ontogeny. Moreover, they acknowledge Daniele Armaleo and Basil Britto for providing access to the organelle genomes of *Cladonia grayi* & *Asterochloris glomerate*, and Olafur S. Andresson for sharing the *Lobaria pulmonaria* data.

