## Supplementary Material for "What is in a lichen? A metagenomic approach to reconstruct the holo-genome of *Umbilicaria pustulata*"

### Supplementary Materials

Supplementary Table S7 – Non-detected $LCA_{Lec}$ genes in the *U. pustulata* data .9
Supplementary Figure S19 – Gene structure of *U. pustulata* (mycobiont) *nad6* 27
Supplementary Figure S20 – *nad3* locus of *U. pustulata* (mycobiont) ....28
Supplementary Figure S21 – Relocation of the chloroplast *rps4* gene to the nuclear genome of *Trebouxia sp.*....28

### Supplementary text

#### Assembly of the lichen holo-genome

A comprehensive reconstruction of genome sequences from metagenomic sequencing data with varying coverages is typically not achievable with the use of only a single assembler (Chakraborty et al. 2016). Here, we pursued a stepwise assembly strategy to target the different genomes in the lichen holo-genome. An overview of the complete processing and assembly workflow is shown in Figure S1.

#### High Coverage Fractions

We used *FALCON v0.2.1* (Chin et al. 2016) to assemble the high coverage fraction of the *L. pustulata* metagenome, using the unprocessed PacBio data. For the initial mapping and the pre-assembly, we used a *length_cutoff* of 3,500. For the preassembly the *--min_idt 0.70 --min_cov 3 --local_match_count_threshold 2 --max_n_read 200* flags were used. Overlap filtering was performed with the following parameter settings: *--max_diff 100 --max_cov 100 --min_cov 2 --bestn 10*.

#### Low Coverage Fractions

To target the genomic fractions with a lower coverage in the metagenomic reads, we employed two different assembly strategies. We first assembled the PacBio reads that were error-corrected with *ECTools* with the *Celera assembler wgs v8.3rc2* (Berlin et al. 2015) with default parameters. In a parallel approach, we integrated the *ECtools*-corrected *PacBio* reads with data from the two Illumina whole genome shotgun libraries and performed a hybrid-assembly with *SPAdes v3.5.0* (Bankevich et al. 2012)

#### Assembly Merging & Polishing

The *FALCON* assembly was scaffolded using *SSPACE-Long v1.1 (Boetzer and Pirovano 2014)*, using standard parameters. A taxonomic assignment of the resulting scaffolds was done with *DIAMOND v0.6.12.47* (Buchfink, Xie and Huson 2015) & *MEGAN5* (Huson et al. 2011), using a custom database of 120 fungi, 20 plants including four green algae, eight animals, 1471 bacteria and 560 viruses. For the MEGAN analysis, we set the minimum score to 50 and deactivated the low complexity filter for the LCA assignment of *MEGAN*. The same workflow was used to taxonomically classify the contigs of the *SPAdes* and the *Celera* assembly into bacterial, algal and fungal fractions.

The fungal *SPAdes* contigs of a length larger than 3 Kbp were used to additionally scaffold the fungal *FALCON* contigs. For the algal genome, we merged contigs of all three assemblies assigned to the *Viridiplantae* into a single algal assembly using *minimus2 (Treangen et al. 2011)*. This merged assembly was then subjected to a final scaffolding step using *SSPACE-Long*. The bacterial contigs from the three assemblies were merged following the same procedure as for the algal sequences.

For assembly polishing, we mapped the preprocessed metagenomic *Illumina* reads to the three assemblies with *bowtie2* (Langmead and Salzberg 2012). This data was then used as input for *Pilon v1.15* (Walker et al. 2014). For polishing of fungal assembly as well as for the organellar genome assemblies, we considered only reads from the mate pair library representing a lichen thallus from the same geographic region as that which was used for the PacBio sequencing. For the lower-coverage bacterial and algal assembly both read- and mate-pairs were used, as the *Illumina* coverage would have not sufficed otherwise.

#### Genome Annotation

##### Fungal Genome Annotation

We annotated genes in the fungal genome with *funannotate* (https://github.com/nextgenusfs/funannotate). For training the pipeline we used both the RNAseq data of *L. pustulata* as well as the proteomes of the lichenized fungi *Xanthoria parietina* and *Cladonia grayi*. In a first step we aligned the processed RNAseq data to the assembled fungal genome using *HISAT2* (Kim, Langmead and Salzberg 2015), using *–max-intronlen 3000* and otherwise standard parameters. In a second step we assembled the processed RNAseq data with *Trinity release 2013-11-10* (Grabherr et al. 2011) using the *de novo* mode with *–jaccard_clip –normalize_reads* as well as the *reference-guided* mode, using the assembly of *L. pustulata* as the reference. The joint output of *Trinity* was then used to run *TransDecoder* through the *PASA* pipeline (Haas et al. 2008), to identify transcript assemblies on the reference assembly. The RNAseq mapping by *HISAT2*, the joint *Trinity* assemblies as well as the *PASA* results and the proteomes of *X. parietina* and *C. grayi* were finally used for running the gene prediction through *funannotate*. The same workflow was used to annotate the publicly available but unannotated genome of *Umbilicaria muehlenbergii* and the draft genome of *Lasallia hispanica*.

#### Algal Genome Annotation

The annotation of the *Trebouxia sp*. was done using *Maker* in a two-step iterative approach. In the first step *Maker* was run with gene predictions made by the *CEGMA* (Parra, Bradnam and Korf 2007) that were converted to SNAP HMMs, the assembled RNAseq data, the proteome of *Asterochloris sp.* and *GeneMark* (Besemer and Borodovsky 2005). The resulting gene predictions were used to train *AUGUSTUS v3.1* (Stanke et al. 2006) and *SNAP v2006-07-28 (Korf 2004)*. These new models, along with *GeneMark* and the RNA/Protein evidences were used for a second round of gene predictions with *Maker*, using more stringent settings.

#### Absence of lecanoromycete core genes

Evolutionary events, i.e. gene loss, or artefacts during the reconstruction of gene sets from draft genomes, determine jointly the set of evolutionarily old genes that are deemed absent from a the genome sequence of a newly sequence organism (Deutekom et al. 2019). To assess the impact of reconstruction artefacts beyond what can be achieved with a standard BUSCO analysis, we investigated the presence-absence pattern of genes that can be traced back to the last common ancestor of the Lecanoromycetes ($LCA_{Lec}$). OMA standalone (Altenhoff et al. 2018) was used to compute hierarchical orthologous groups (HOGs) for five lecanoromycete taxa: *L. pustulata*, *Umbilicaria muehlenbergii*, *Usnea florida*, *Cladonia grayi*, and *Xanthoria parietina*. All five species were represented in 4,607 HOGs. We subsequently concentrated on 1,402 HOGs with one taxon missing. We performed a three-staged procedure to search for the missing ortholog in the gene set and, if necessary, in the genome sequence and the transcriptome of the respective species. In stage one, the orthologous group served as input for the more inclusive targeted ortholog search tool *HaMStR* (Ebersberger, Strauss and von Haeseler 2009b). Notably, the lengths of the orthologs found via *HaMStR* but missed by OMA differ substantially from that of the other sequences in the HOG (supplementary figure S8, Supplementary Material online). These differences may indicate artificial gene fusions in the process of genome annotation. Since intergenic spaces are typically small in fungi, this is a common artefact in fungal genome annotation (Testa et al. 2015). Orthologous groups that could not be completed were promoted to the second stage, where they served as input for an *exonerate* search (Slater and Birney 2005) in the genome sequence. For *exonerate* we aligned all proteins present in a given orthologous group against the genomes of all five species. We assumed a gene to be truly absent if the alignment scores against the genome of the species, where the gene is missing, are significantly worse (Mann-Whitney-Wilcoxon test, p<= 0.05) when compared to the alignment score distribution observed when aligning against the genomes where the ortholog has been found. This provides evidence that up to $2/3_{rd}$ (*U. muehlenbergii*) of the missing $LCA_{Lec}$ genes have been overlooked during the gene annotation, as *exonerate* identifies candidate regions where no gene has been annotated. In the third stage, we searched for the missing genes in the assembled transcript data with *HaMStR* using the option -est (Ebersberger, Strauss and von Haeseler 2009a). The additional orthologs identified here most likely belong to genes that reside in assembly gaps. The results of the searches are summarized in supplementary table S7, Supplementary Material online.

#### Supplementary tables

##### Supplementary Table S1 – Composition of the custom database for the taxonomic assignment of contigs/scaffolds

See file *SupplTabS1_6_8_9.xslx*

##### Supplementary Table S2 – qPCR analysis of the fungal to algal cell ratio in the thalli of *Lasallia pustulata*

| | | Copy number per replicate | | | | |
|---|---|---|---|---|---|---|
| **Thallus** | | **#1** | **#2** | **#3** | **Mean** | **Fungal:Algal ratio** |
| **1** | Fungal | 24,618 | 28,317 | 27,952 | 26,963 | 14 |
| | Algal | 1,868 | 1,972 | 1,928 | 1,928 | |
| **2** | Fungal | 27,375 | 28,196 | 30,826 | 28,799 | 13 |
| | Algal | 1,994 | 2,368 | 2,458 | 2,274 | |
| **3** | Fungal | 36,805 | 39,654 | 40,730 | 39,064 | 14 |
| | Algal | 2,581 | 2,863 | 2,745 | 2,730 | |
| **4** | Fungal | 36,769 | 37,257 | 38,935 | 37,654 | 24 |
| | Algal | 1,478 | 1,531 | 1,694 | 1,568 | |

#### Supplementary Table S3 – Metrics of the metagenome assembly

| Assembly method | Taxonomic classification | Number of Scaffolds | Total length (Mbp) | N50 (bp) |
|---|---|---|---|---|
| **FALCON** | All | 2,343 | 62 | 322,812 |
| | Fungal | 120 | 32 | 550,723 |
| | Algal | 709 | 9 | 17,208 |
| | Bacterial | 790 | 15 | 56,167 |
| **SPAdes** | All | 21,900 | 123 | 224,806 |
| | Fungal | 5,736 | 35 | 159,267 |
| | Algal | 257 | 47 | 461,016 |
| | Bacterial | 1,193 | 26 | 91,450 |
| **Celera** | All | 22,216 | 216 | 11,162 |
| | Fungal | 12,230 | 113 | 10,395 |
| | Algal | 3,557 | 52 | 16,617 |
| | Bacterial | 2,804 | 17 | 7,798 |
| **Final#** | Fungal | 43 | 33 | 1,808,250 |
| | Algal | 225 | 53 | 848,255 |
| | Bacterial | 499 | 35 | 250,871 |

### Final assembly steps comprise the merging of contigs from different assembly strategies assigned to the same taxonomic group with Minumus, followed by a final scaffolding step.

#### Supplementary Table S4 Coverage ratios in the lichen holo-genome

| | PacBio | PE | MP | Pool1 | Pool2 | Pool3 | Pool4 | Pool5 | Pool6 | Average |
|---|---|---|---|---|---|---|---|---|---|---|
| ***Trebouxia sp.* nuclear#** | 1.00 | 1.00 | 1.00 | 1.00 | 1.00 | 1.00 | 1.00 | 1.00 | 1.00 | 1.0 |
| ***Trebouxia sp.* cp** | 23.79 | 8.00 | 8.7 | 21.6 | 14.3 | 18.8 | 19.9 | 16.7 | 13.6 | 16.2 |
| ***Trebouxia sp.* mt** | 21.3 | 9.1 | 4.2 | 25.6 | 16.7 | 21.5 | 19.2 | 14.3 | 17.4 | 16.6 |
| ***U. pustulata* nuclear** | 13.8 | 11.7 | 17.0 | 19.7 | 19.5 | 16.0 | 24.1 | 28.0 | 29.7 | 19.8 |
| ***U. pustulata* mt** | 329.9 | 229.0 | 239.6 | 261.4 | 196.0 | 228.1 | 435.7 | 379.4 | 322.8 | 287.4 |
| ***Acidobacterium C16*** | 2.5 | 0.4 | 1.0 | 0.5 | 0.7 | 0.6 | 0.6 | 0.8 | 0.7 | 0.9 |
| ***Acidobacterium C35*** | 3.5 | 0.3 | 1.35 | 1.5 | 0.9 | 1.3 | 1.1 | 1.4 | 2.1 | 1.4 |

### All coverages were normalized to the read coverage of *Trebouxia sp*. Data from the *U. pustulata* PoolSeq experiments were obtained from Dal Grande et al. (2017)

#### Supplementary Table S5 – Accession details for the genomes listed in Table 1 (main text)

| Fungal genomes | | Algal genomes | |
|---|---|---|---|
| Species | Accession | Species | Accession |
| *A. radiata* | GCA_002989075.1 | *A. glomerata* | Astpho v2.0 JGI |
| *C. grayi* | Clagr3 v2.0 JGI | *A. protothecoides* | GCA_000733215.1 |
| *C. linearis* | GCA_003521265.1 | *Chlorella* sp. A99 | GCA_003063905.1 |
| *C. macilenta* | AUPP00000000.1 | *C. subellipsoidea* | GCA_000258705.1 |
| *C. metacorallifera* | AXCT00000000.2 | *Helicosporidium sp.* | GCA_000690575.1 |
| *C. rangiferina* | VALI00000000.1 | *T. gelatinosa* | GCA_000818905.1 |
| *C. uncialis* | NAPT00000000.1 | *Trebouxia* sp. TZW2008 | GCA_002118135.1 |
| *E. prunastri* | NKYR00000000.1 | | |
| *E. pusillum* | GCA_000464535.1 | | |
| *G. flavorubescens* | AUPK00000000.1 | | |
| *U. hispanica* | GCA_003254425.1 | | |
| *L. pulmonaria* | JGI Lobpul1 v1.0 | | |
| *P. furfuracea* | GCA_003184345.1 | | |
| *R. intermedia* | PEKF00000000.1 | | |
| *R. peruviana* | GCA_001956345.1 | | |
| *U. florida* | JGI Usnflo1 v1.0 | | |
| *U. muehlenbergii* | GCA_000611775.1 | | |
| *X. parietina* | Xanpa2 v1.1 JGI | | |

#### Supplementary Table S6 – Diamond results for the genes proposed as HGT candidates in the *U. pustulata* genome

See file *SupplTabS1_6_8_9.xslx*

#### Supplementary Table S7 – Non-detected $LCA_{Lec}$ genes in the *U. pustulata* data

| Taxon | Annotated gene set (*HaMStR*)§ | *Assembly (Exonerate)* § | *Transcripts* (HaMStR) § |
|---|---|---|---|
| *U. pustulata* | 142 | 76 | 33 |
| *U. muehlenbergii* | 126 | 36 | *NA** |
| *C. grayi* | 157 | 58 | 52 |
| *U. florida* | 90 | 55 | 45 |
| *X. parietina* | 203 | 159 | 90 |

§Numbers of LCALec genes exclusively missing in the respective species. Only the genes found missing in a previous analysis served as input for the next search
*no RNAseq data available

**Supplementary Table S8 – $LCA_{Lec}$ genes that appear genuinely absent from the *U. pustulata* genome before and after manual curation**

See file *SupplTabS1_6_8_9.xslx*

**Supplementary Table S9 – Representation of core genes in the mitochondrial genome of *U. pustulata***

See file *SupplTabS1_6_8_9.xslx*

### Supplementary Figures

#### Supplementary Figure S1 – The assembly workflow

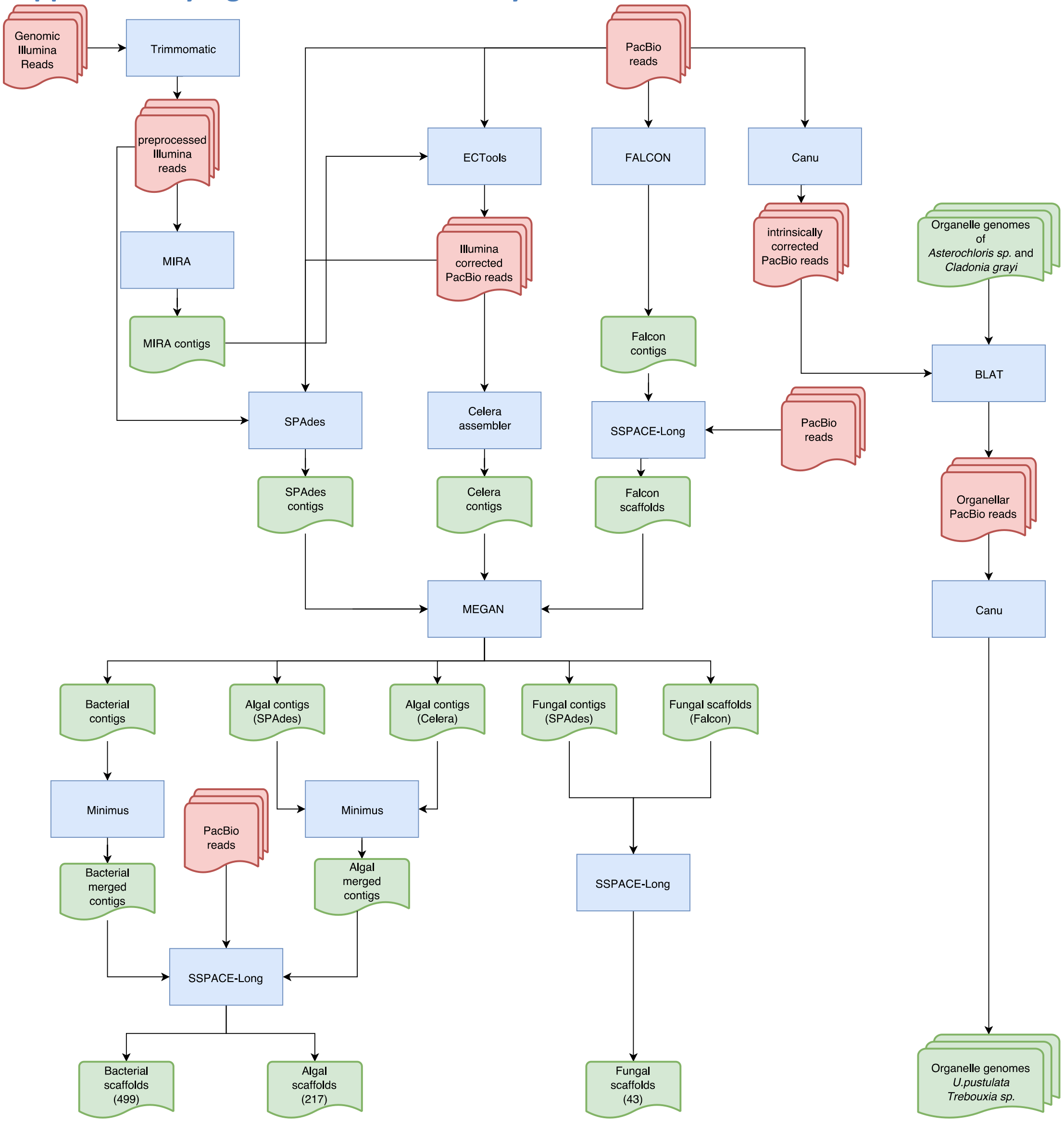

**Figure S1 – The assembly workflow**. Color code: red – read data; green – assembled contigs/scaffolds; blue – software. See supplementary text, Supplementary Material online, for further details.

#### Supplementary Figure S2 – The mitochondrial genome of the fungus *U. pustulata*

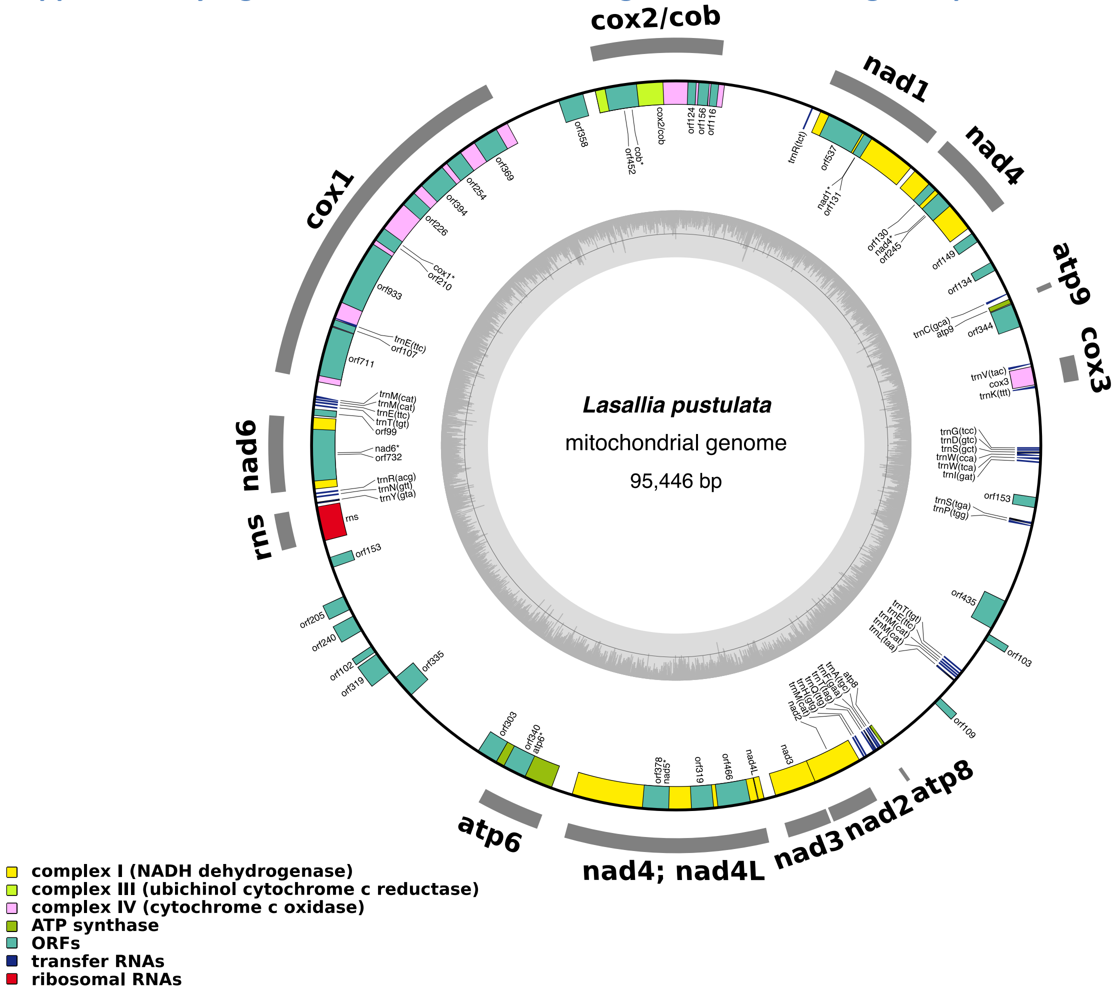

**Supplementary Figure S2 – The mitochondrial genome of the fungus *U. pustulata*.** The mitochondrial genome has a total length of 95 kb. It harbors 15 protein coding genes, a small subunit rRNA gene, 33 additional ORFs, and 31 tRNA genes. The longest gene, cox1, comprises 8 exons and spans 14 kb. cox2 and cob (cytochrome oxidase b) are fused into a single gene.

#### Supplementary Figure S3 – The mitochondrial genome of *Trebouxia sp.*

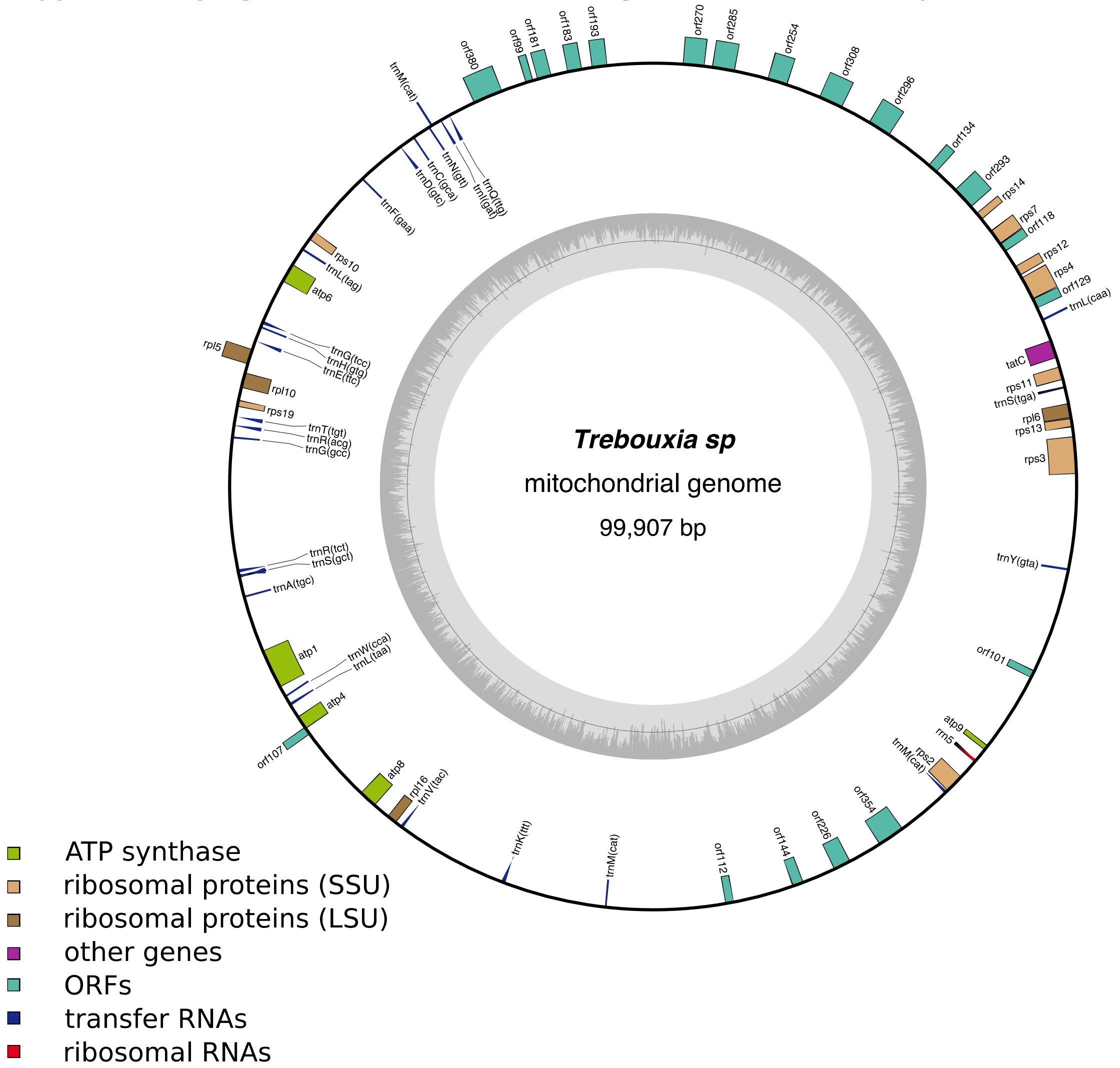

**Supplementary Figure S3 – The mitochondrial genome of the alga *Trebouxia sp.*.** The mitochondrial genome has a total length of 99,9 kb. It harbors 32 protein coding genes, an rRNA gene, 20 additional ORFs, and 26 tRNA genes.

#### Supplementary Figure S4 – The chloroplast genome of *Trebouxia sp.*

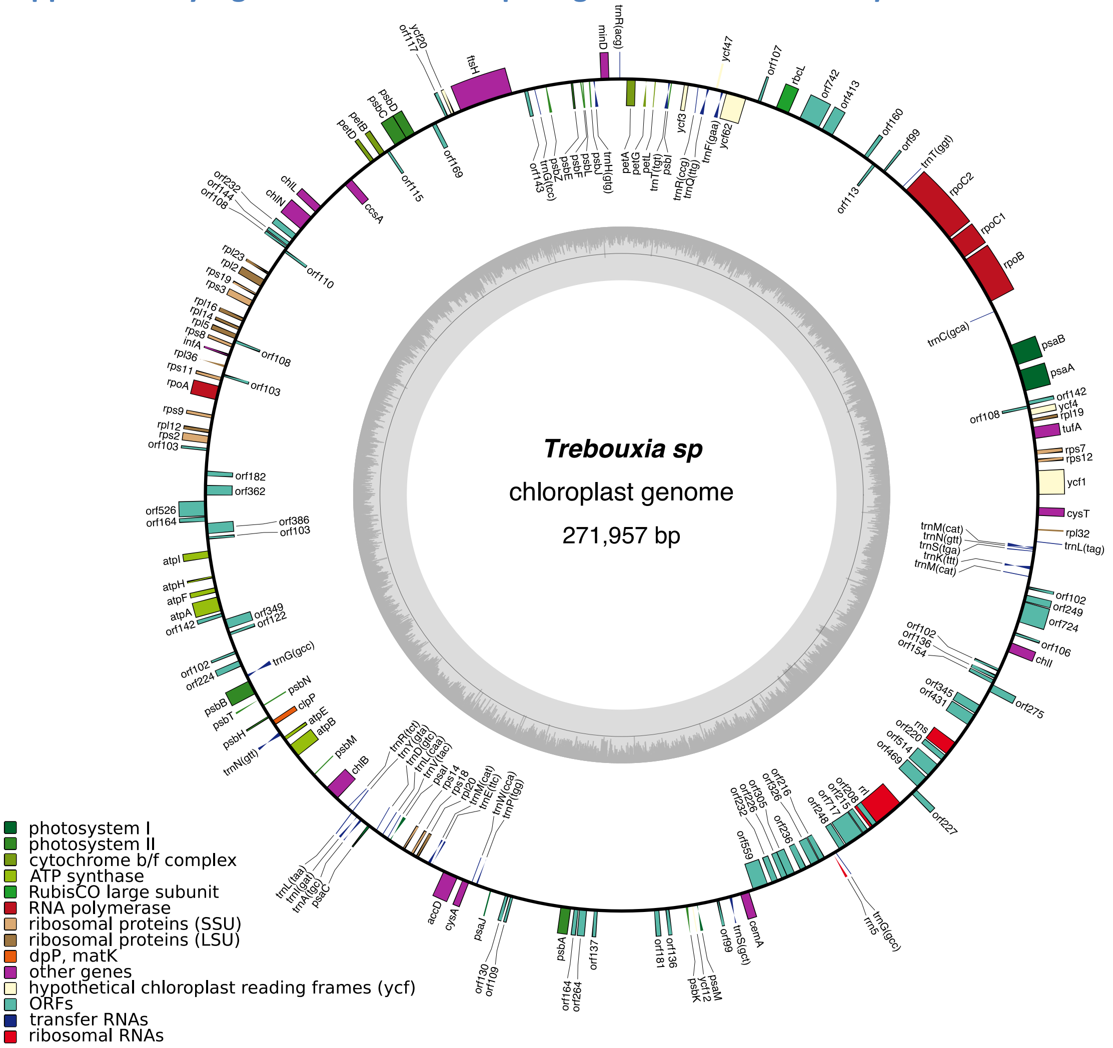

**Supplementary Figure S4 – The chloroplast genome of the alga *Trebouxia sp.*.** The chloroplast genome has a total length of 271,96 kb. It harbors 78 protein coding genes, three rRNA genes, 52 additional ORFs, and 31 tRNA genes.

#### Supplementary Figure S5 – Composition of the *U. pustulata* microbiome on a species level resolution

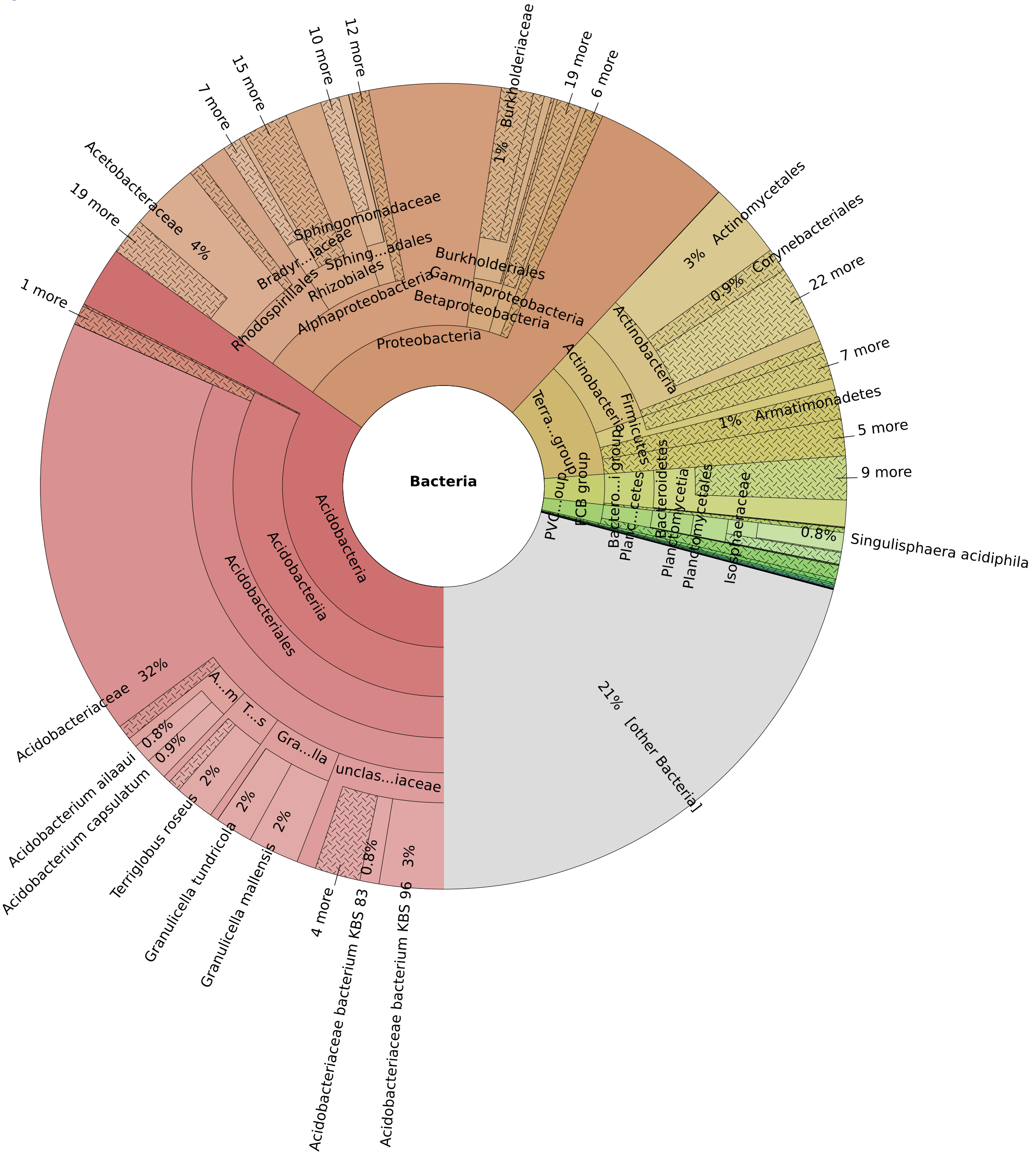

**Supplementary Figure S5 – Composition of the *U. pustulata* microbiome on a species level resolution.** Reads from the two Illumina whole genome shotgun libraries and the PacBio reads were pooled and taxonomically assigned to the species level with MEGAN (Huson et al. 2016). *Acidobacteriaceae* are known to be tolerant to changes in hydration (Ward et al. 2009), and the two *Granulicella* species, *G. tundricola* and *G. mallensis*, are dominant members of the soil bacterial community in the arctic alpine tundra (Rawat et al. 2014, Rawat et al. 2013). They have been described as versatile heterotrophs that hydrolyze a suite of sugars and complex polysaccharides including plant-based carbon polymers. Thus, *Acidobacteriaceae* in general, and the two *Granulicella* species in particular, are well adapted to live in the same environment as *U. pustulata*, which preferentially grows on rocks with frequent cycles of drying and rewetting (Sletvold and Hestmark 1997, Dal Grande et al. 2017).

#### Supplementary Figure S6 – The impact of different taxonomic assignment strategies on taxon abundance estimates

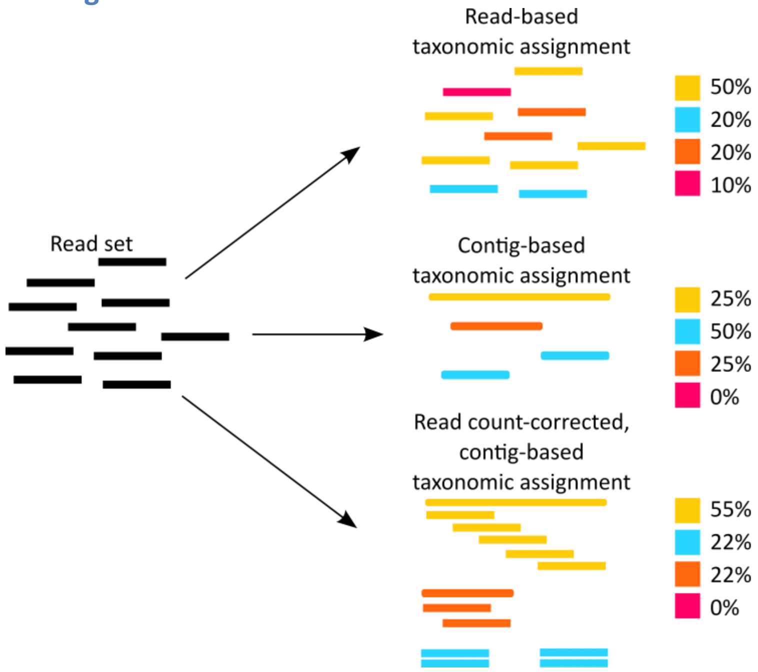

**Supplementary Figure S6 - Different taxonomic assignment strategies, and their impact on taxon abundance estimates**. Ideally, the number of reads representing an y particular taxon in a metagenome is proportional to its average genome copy number (but see (Nayfach and Pollard 2016)). Given that all reads can be accurately taxonomically classified, read-based analyses (top) result in accurate abundance estimates. A taxonomic assignment on the contig-level (middle) counts long contigs, consisting of many reads, only once. As a consequence, the abundance of the yellow taxon is underestimated, while abundance estimate for the orange and the blue taxon, respectively, increase. Note, reads from the red taxon remained unassembled, and thus this taxon is missed. A hybrid approach (bottom) performs the taxonomic assignment on the contig level. Taxon abundance estimates are then based on the reads mapping to the contigs. While the use of coverage information can correct for the bias introduced by the contig-based taxonomic assignment, taxa whose reads could not be assembled, will still be missed.

**Supplementary Figure S7 – Taxonomic composition of the *U. pustulata* microbiome based on the assembled reads, without (A) and with (B) taking the read coverage into account.**

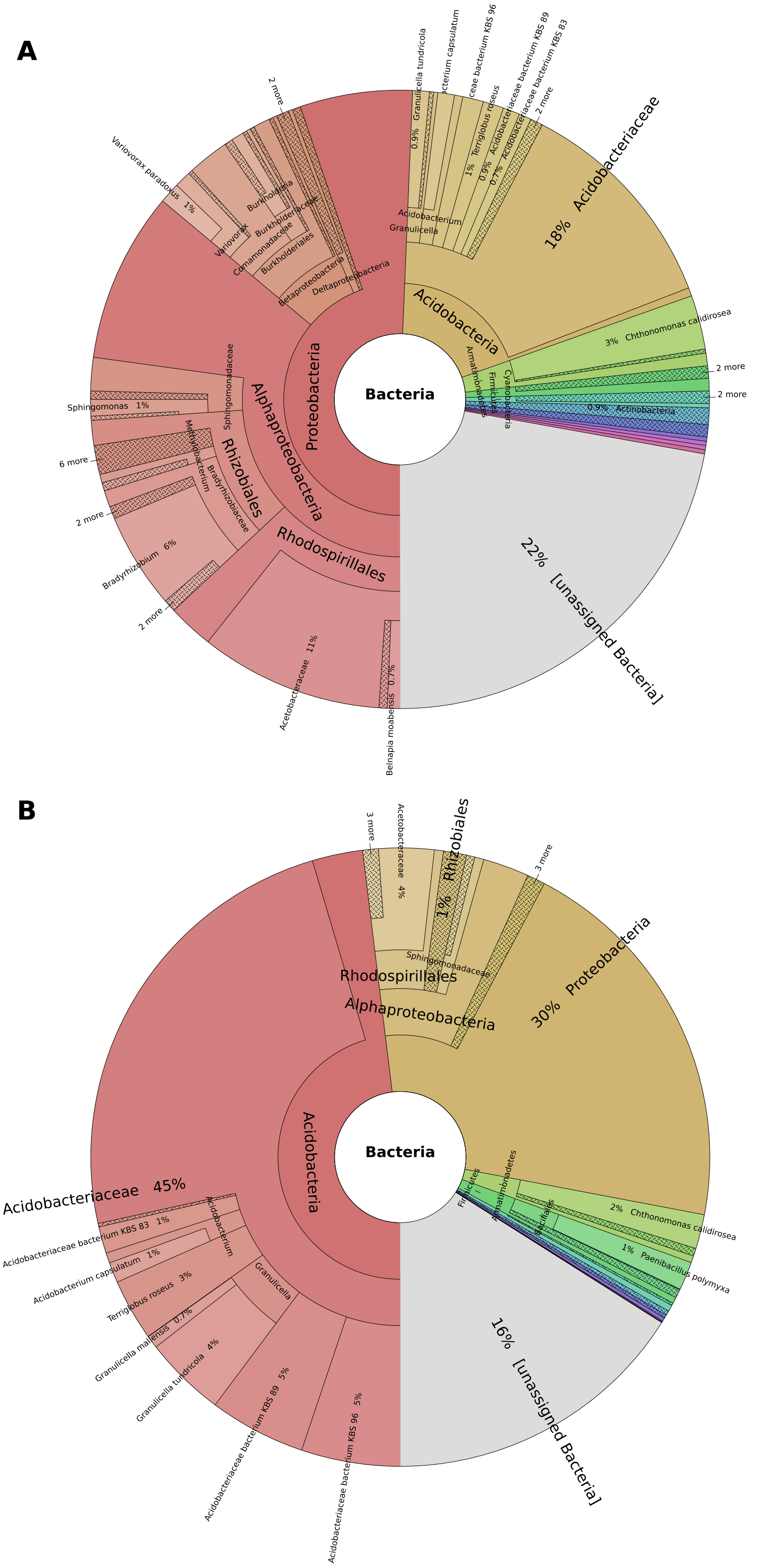

**Supplementary Figure S7** – Taxonomic composition of the *U. pustulata* microbiome based on the assembled contigs. The color code follows taxon abundance. We assessed the abundance of the bacterial taxa using the assembled contigs instead of the individual reads **(A)**. This shifted the estimated abundances of the individual taxa in favor of the *Proteobacteria*, and the contribution of *Rhizobiales* almost tripled to 11%. In turn, the abundance estimate for the Acidobacteriaceae decreased to 18%. When we corrected the estimates by using the number of mapped reads to each scaffold for the basis of the classification, the abundance of the *Rhizobiales* droped to 1%, while that of the *Acidobacteriaceae* grew to 45%. The difference to the abundance estimates on the read level (*see* Figure 2 in the main text) are due to reads that could not be assembled, and hence were not considered in the contig-based analysis.

#### Supplementary Figure S8 – Length difference between orthologs missed by OMA and detected by HaMStR

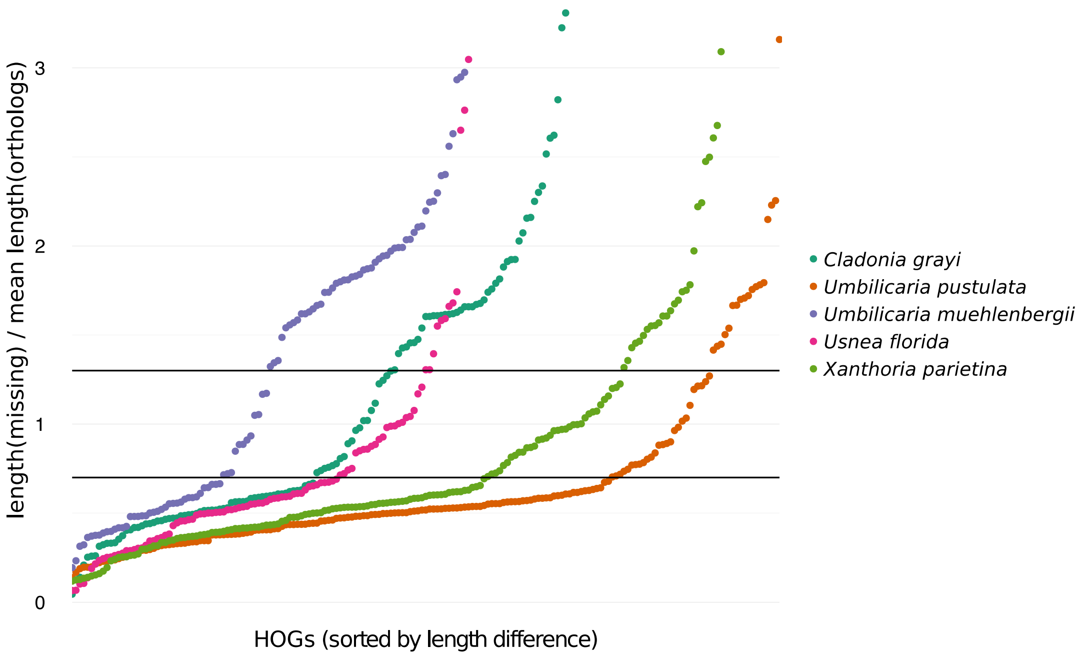

**Supplementary Figure S8 – Length difference between orthologs missed by OMA and detected by HaMStR.** To assess the reason why individual genes have been missed by OMA but were predicted as orthologs by HaMStR, we computed the ratio between the sequence lengths of the corresponding *U. pustulata* proteins and the average length of the proteins in the orthologous group. The horizontal bars give the upper and lower bound of the length cut-off filter used in the ortholog identification of OMA. This reveals that most of the orthologs missed by OMA exceed the length cutoff implemented into the OMA algorithm, most likely due to the artificial fusion or fission of genes in the course of gene annotation.

#### Supplementary Figure S9 – Curation of the frameshift mutations in the DHFR locus – Deletion 1

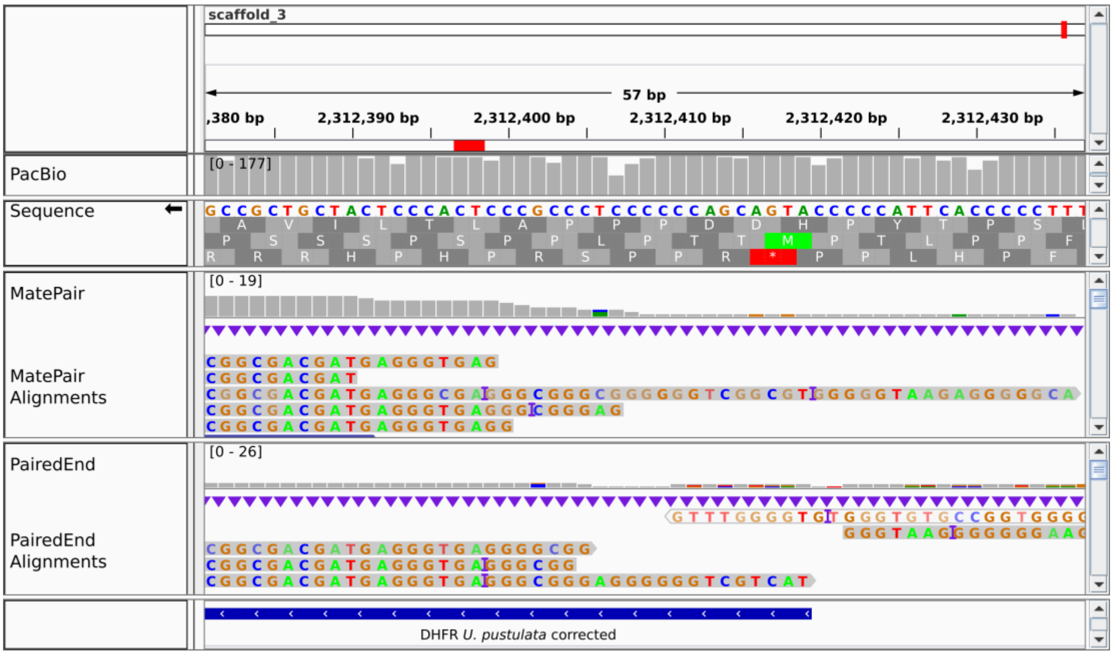

**Supplementary Figure S9 – Curation of the frameshift mutations in the DHFR locus – Deletion 1.** The red bar indicates the position of the frameshift. “Sequence” provides the consensus information of the assembly (reverse complemented) together with the translation in 3 reading frames. “PacBio”, “MatePair”, and “PairedEnd” tracks provide the read coverage information for this window. “MatePair Alignment” and “PairedEnd Alignment” show the mapped reads from the respective libraries (not reverse complemented). The purple bar indicates a deletion in the read relative to the reference. All informative PairedEnd and MatePair reads support four rather than just the three Cs represented in the reference sequences.

#### Supplementary Figure S10 – Curation of the frameshift mutations in the DHFR locus – Deletion 2

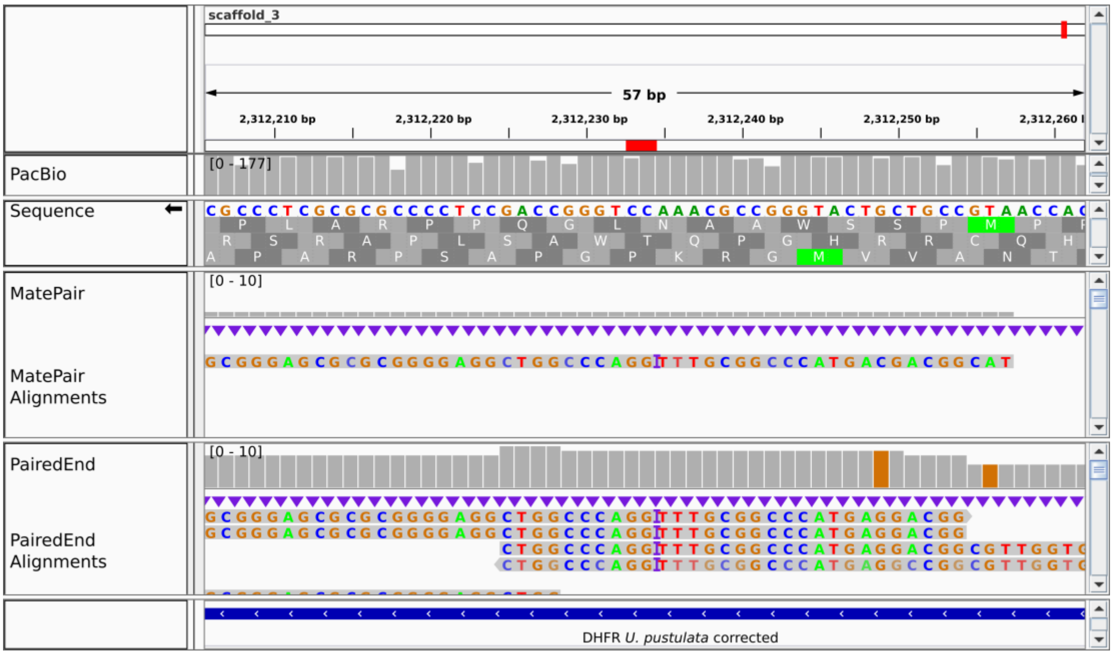

**Supplementary Figure S10 – Curation of the frameshift mutations in the DHFR locus – Deletion 1.** The red bar indicates the position of the frameshift. “Sequence” provides the consensus information of the assembly (reverse complemented) together with the translation in 3 reading frames. “PacBio”, “MatePair”, and “PairedEnd” tracks provide the read coverage information for this window. “MatePair Alignment” and “PairedEnd Alignment” show the mapped reads from the respective libraries (not reverse complemented). The purple bar indicates a deletion in the read relative to the reference. All informative PairedEnd and MatePair reads support four rather than just the three As represented in the reference sequences.

#### Supplementary Figure S11 – Curation of the frameshift mutations in the DHFR locus – Deletion 3

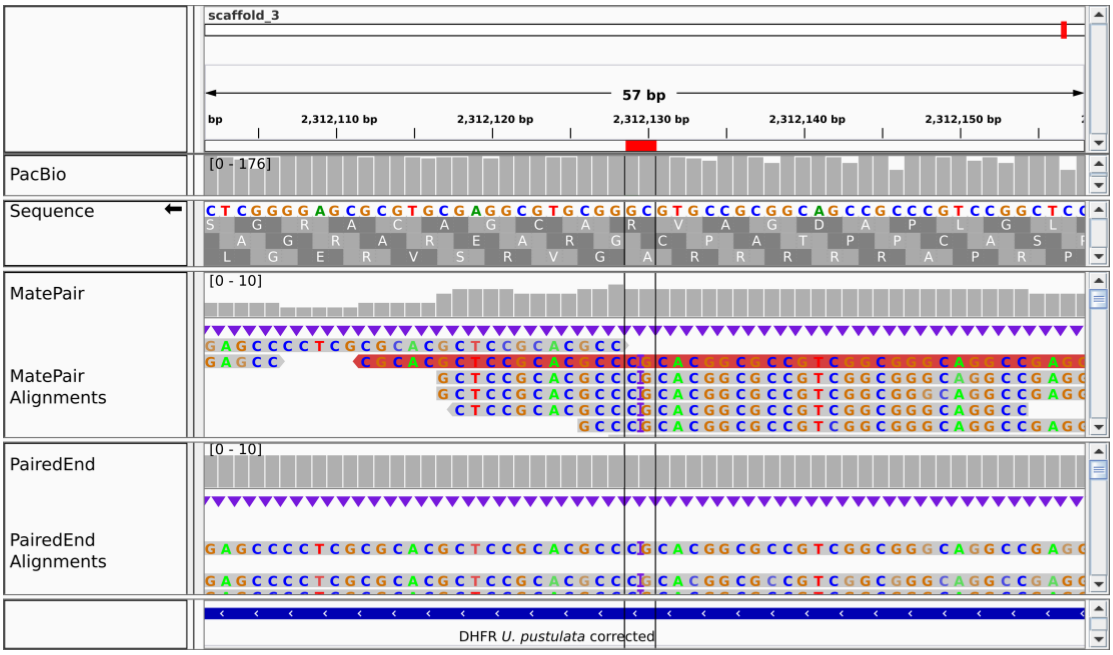

**Supplementary Figure S11 – Curation of the frameshift mutations in the DHFR locus – Deletion 3.** The red bar indicates the position of the frameshift. “Sequence” provides the consensus information of the assembly (reverse complemented) together with the translation in 3 reading frames. “PacBio”, “MatePair”, and “PairedEnd” tracks provide the read coverage information for this window. “MatePair Alignment” and “PairedEnd Alignment” show the mapped reads from the respective libraries (not reverse complemented). The purple bar indicates a deletion in the read relative to the reference. All informative PairedEnd and MatePair reads support an additional A between the consecutive CG represented in the reference sequences.

#### Supplementary Figure S12 – Curation of the frameshift mutations in the DHFR locus – Deletion 4

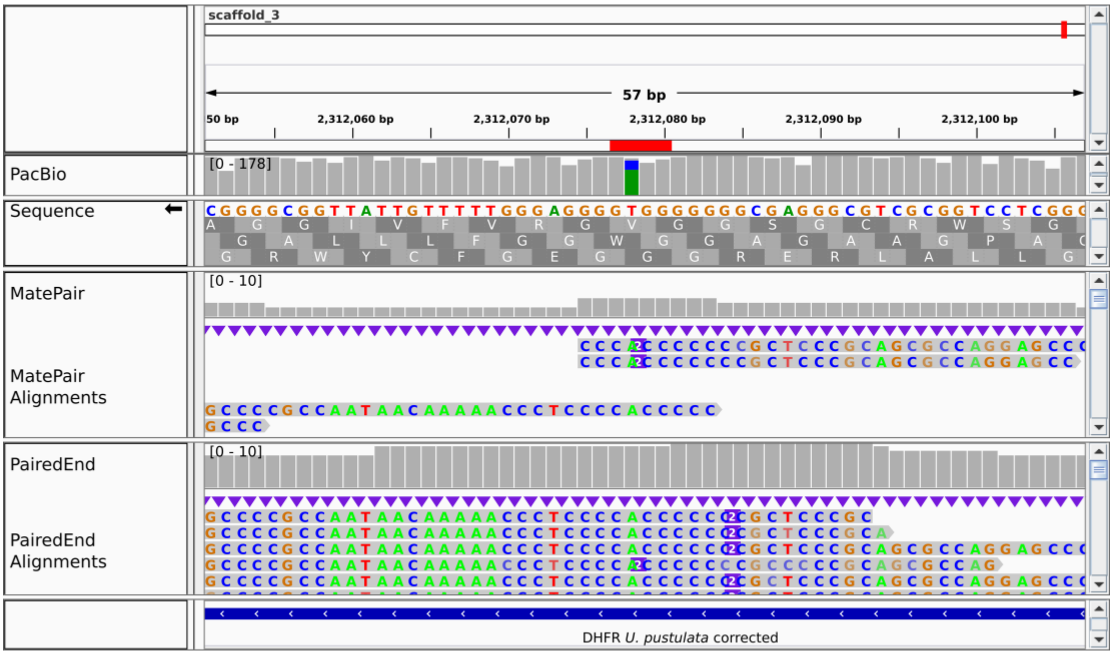

**Supplementary Figure S12 – Curation of the frameshift mutations in the DHFR locus – Deletion 4.** The red bar indicates the position of the frameshift. “Sequence” provides the consensus information of the assembly (reverse complemented) together with the translation in 3 reading frames. “PacBio”, “MatePair”, and “PairedEnd” tracks provide the read coverage information for this window. “MatePair Alignment” and “PairedEnd Alignment” show the mapped reads from the respective libraries (not reverse complemented). The purple bar indicates a deletion in the read relative to the reference. All informative PairedEnd and MatePair reads support the existence of two additional Gs relative to the reference sequences.

#### Supplementary Figure S13 – Curation of the frameshift mutations in the DHFR locus – Deletion 5

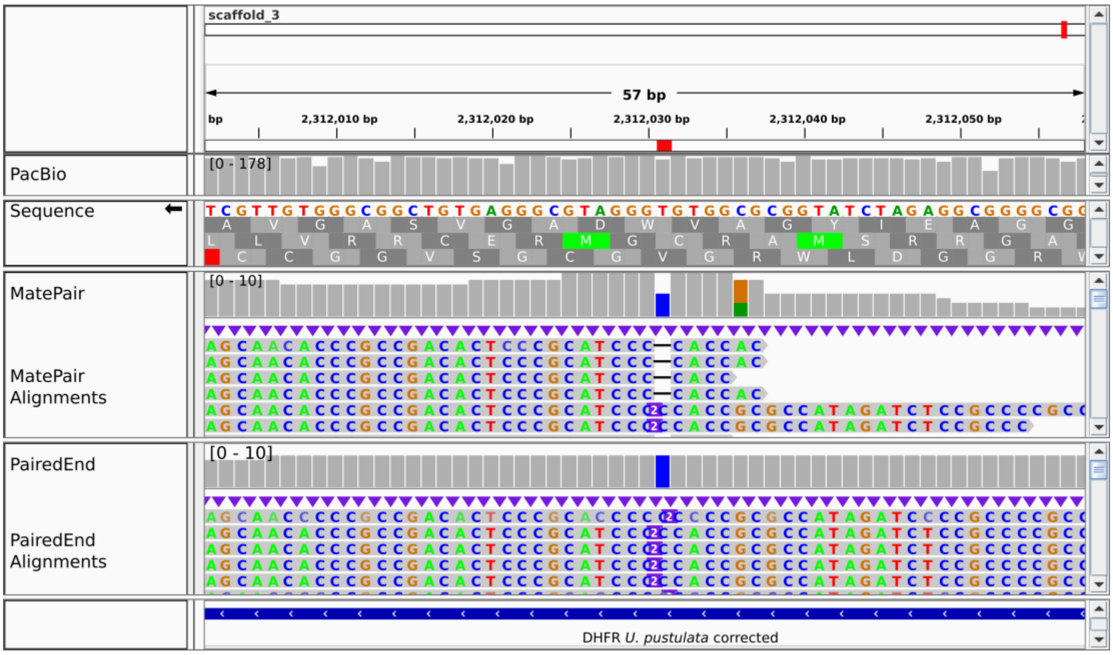

**Supplementary Figure S13 – Curation of the frameshift mutations in the DHFR locus – Deletion 5.** The red bar indicates the position of the frameshift. "Sequence" provides the consensus information of the assembly (reverse complemented) together with the translation in 3 reading frames. "PacBio", "MatePair", and "PairedEnd" tracks provide the read coverage information for this window. "MatePair Alignment" and "PairedEnd Alignment" show the mapped reads from the respective libraries (not reverse complemented). The purple bar indicates a deletion in the read relative to the reference. All informative PairedEnd and MatePair reads support the existence of an additional TG nucleotide plus the exchange of a T to G relative to the reference sequences.

#### Supplementary Figure S14 – Curation of the frameshift mutations in the DHFR locus – Deletion 6

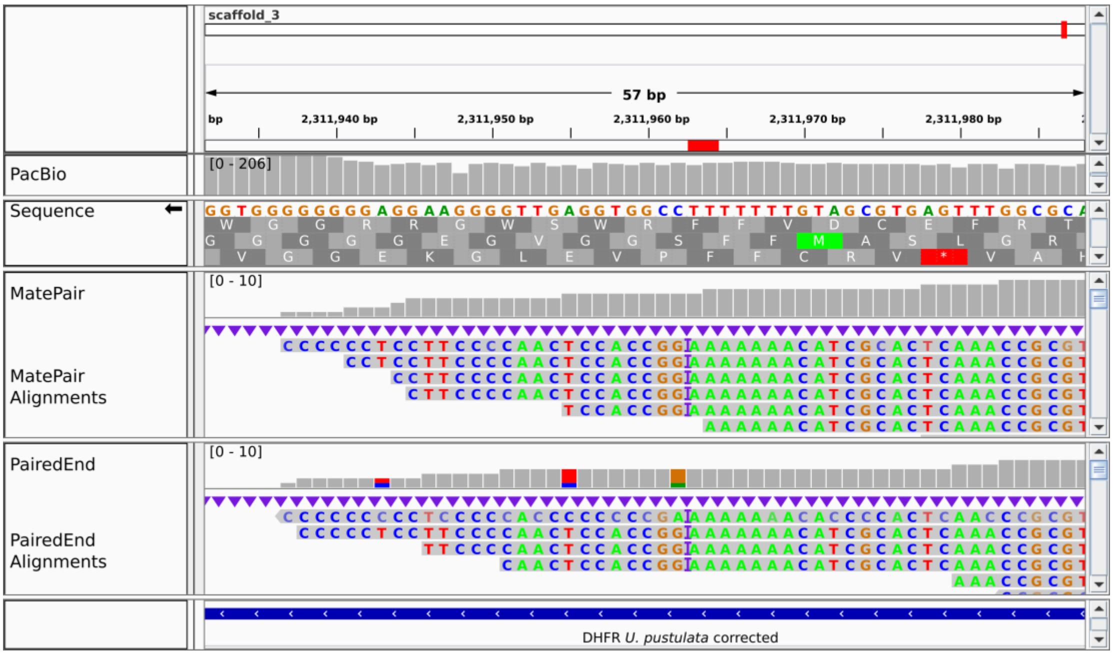

**Supplementary Figure S14 – Curation of the frameshift mutations in the DHFR locus – Deletion 6.** The red bar indicates the position of the frameshift. "Sequence" provides the consensus information of the assembly (reverse complemented) together with the translation in 3 reading frames. "PacBio", "MatePair", and "PairedEnd" tracks provide the read coverage information for this window. "MatePair Alignment" and "PairedEnd Alignment" show the mapped reads from the respective libraries (not reverse complemented). The purple bar indicates a deletion in the read relative to the reference. All informative PairedEnd and MatePair reads support the existence of an additional T relative to the reference sequences.

#### Supplementary Figure S15 – Curation of the frameshift mutations in the DHFR locus – Deletion 7

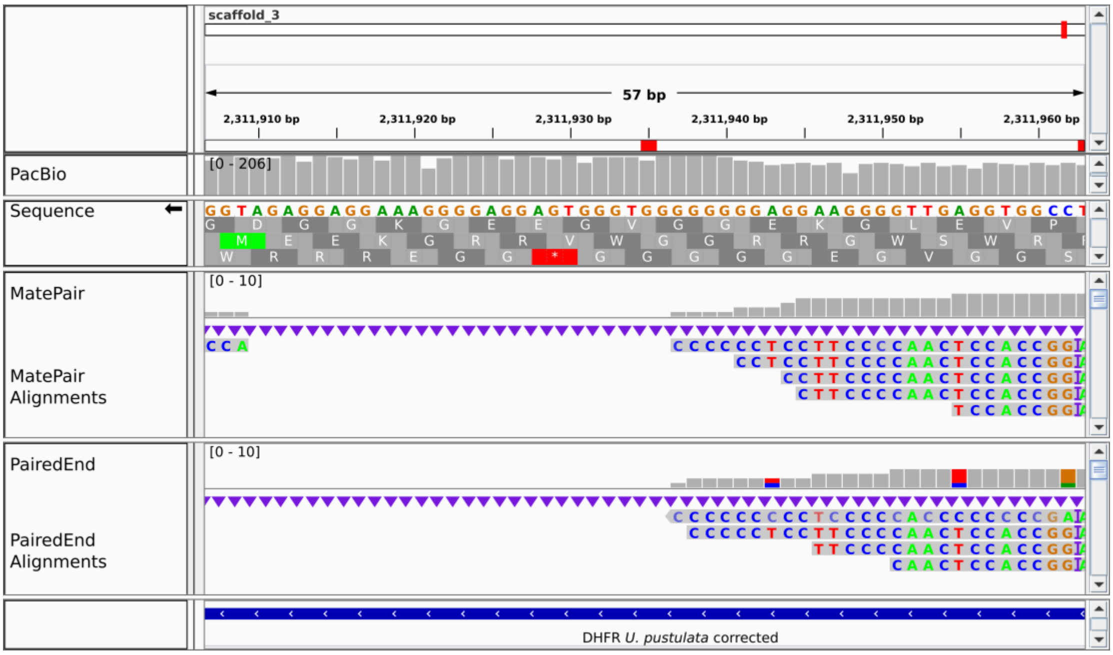

**Supplementary Figure S15 – Curation of the frameshift mutations in the DHFR locus – Deletion 7.** The red bar indicates the position of the frameshift. “Sequence” provides the consensus information of the assembly (reverse complemented) together with the translation in 3 reading frames. “PacBio”, “MatePair”, and “PairedEnd” tracks provide the read coverage information for this window. “MatePair Alignment” and “PairedEnd Alignment” show the mapped reads from the respective libraries (not reverse complemented). No reads span the deletion causing the frameshift. A comparison to DHFR from XYZ reveals that the extension of the oligo-dG run by two nucleotides rescues the ORF.

#### Supplementary Figure S16 – Curation of the frameshift mutations in the DHFR locus – Deletion 8

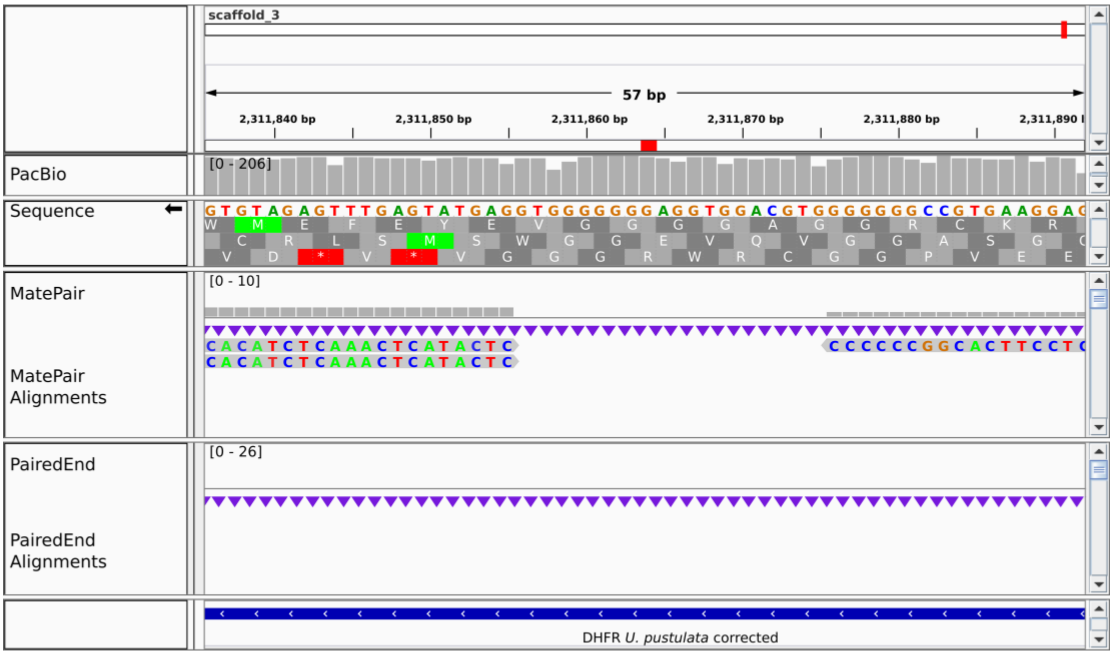

**Supplementary figure S16 – Curation of the frameshift mutations in the DHFR locus – Deletion 8.** The red bar indicates the position of the frameshift. "Sequence" provides the consensus information of the assembly (reverse complemented) together with the translation in 3 reading frames. "PacBio", "MatePair", and "PairedEnd" tracks provide the read coverage information for this window. "MatePair Alignment" and "PairedEnd Alignment" show the mapped reads from the respective libraries (not reverse complemented). No reads span the deletion causing the frameshift. A comparison to DHFR from *U. hispanica* reveals that the extension of the oligo-dG run by two nucleotides rescues the ORF.

#### Supplementary Figure S17 – BlastN alignment of the edited DFHR ORF against the *U. pustulata* reference genome sequence

```
> scaffold_3
Length=2390932

 Score = 1092 bits (591),  Expect = 0.0
 Identities = 621/633 (98%), Gaps = 12/633 (2%)
 Strand=Plus/Minus

Edt.    1         ATGACGACCCCCCTCCCGCCCCTCACCCTCATCGTCGCCGCCACCCACCCGACCCTAGGC  60
                  ||||||||||||||||||||| ||||||||||||||||||||||||||||||||||||||
Orig.   2312419   ATGACGACCCCCCTCCCGCCC-TCACCCTCATCGTCGCCGCCACCCACCCGACCCTAGGC 2312361

Edt.    61        ATCGGCCTCCGCTCCACCCTCCCCTGGCCGCACCTCCCGCGCGAACTCGCCTACTTCGCG  120
                  ||||||||||||||||||||||||||||||||||||||||||||||||||||||||||||
Orig.   2312360   ATCGGCCTCCGCTCCACCCTCCCCTGGCCGCACCTCCCGCGCGAACTCGCCTACTTCGCG  2312301

Edt.    121       CGCACCACGACGCGccccccccccGCCCCACCCAACCGCCACCAATGCCGTCGTCATGGGC  180
                  ||||||||||||||||||||||||||||||||||||||||||||||||||||||||||||
Orig.   2312300   CGCACCACGACGCGCCCCCCCCCCGCCCCACCCAACCGCCACCAATGCCGTCGTCATGGGC  2312241

Edt.    181       CGCAAAACCTGGGCCAGCCTCCCCGCGCGCTCCCGCCCCCTCAAGCGCCGCACCAACGTC  240
                  |||||| |||||||||||||||||||||||||||||||||||||||||||||||||||||
Orig.   2312240   CGCAAA-CCTGGGCCAGCCTCCCCGCGCGCTCCCGCCCCCTCAAGCGCCGCACCAACGTC  2312182

Edt.    241       GTCGTCAGCCGCTCGCCCGACACCCTCGGCCTGCCCGCCGACGGCGCCGTGCAGGGCGTG  300
                  |||||||||||||||||||||||||||||||||||||||||||||||||||| |||||||
Orig.   2312181   GTCGTCAGCCGCTCGCCCGACACCCTCGGCCTGCCCGCCGACGGCGCCGTGC-GGGCGTG  2312123

Edt.    301       CGGAGCGTGCGCGAGGGGCTCCTGGCGCTGCGGGAGCgggggggggTGGGGAGGGTTTTT  360
                  ||||||||||||||||||||||||||||||||||||||||||||  ||||||||||||||
Orig.   2312122   CGGAGCGTGCGCGAGGGGCTCCTGGCGCTGCGGGAGCGGGGGGG--TGGGGAGGGTTTTT  2312065

Edt.    361       GTTATTGGCGGGGCGGAGATCTATGGCGCGGTGGGGGGGATGCGGGAGTGTCGGCGGGTG  420
                  ||||||||||||||||||||||||||||||||| | ||||||||||||||||||||||||
Orig.   2312064   GTTATTGGCGGGGCGGAGATCTATGGCGCGGTG-T-GGGATGCGGGAGTGTCGGCGGGTG  2312007

Edt.    421       TTGCTGACGCGGGTGCGCACGCGGTTTGAGTGCGATGtttttttttCCGGTGGAGTTGGGG  480
                  |||||||||||||||||||||||||||||||||||||||||||| |||||||||||||||
Orig.   2312006   TTGCTGACGCGGGTGCGCACGCGGTTTGAGTGCGATGTTTTTTT-CCGGTGGAGTTGGGG  2311948

Edt.    481       AAGGAggggggggggtgggtgaggaggggaaaggaggagatggatgagtgggtgggggag  540
                  |||||||||||||  |||||||||||||||||||||||||||||||||||||||||||||
Orig.   2311947   AAGGAGGGGGGGG--TGGGTGAGGAGGGGAAAGGAGGAGATGGATGAGTGGGTGGGGGAG  2311890

Edt.    541       gaagtgccgggggggtgcaggtggagggggggggtggagtatgagtttgagatgtgggag  600
                  ||||||||||||||||||||||||||||||||  ||||||||||||||||||||||||||
Orig.   2311889   GAAGTGCCGGGGGGGTGCAGGTGGAGGGGGGG--TGGAGTATGAGTTTGAGATGTGGGAG  2311832

Edt.    601       agggaggggagggaggggaggaggggggggTGA  633
                  |||||||||||||||||||||||||||||||||
Orig.   2311831   AGGGAGGGGAGGGAGGGGAGGAGGGGGGGGTGA  2311799
```

**Supplementary Figure S17 – BlastN alignment of the edited DFHR ORF against the *U. pustulata* reference genome sequence**. The highlighted regions in the edited DHFR ORF ("Edt.") alignment represent the 8 edits that were necessary to recover the full reading frame. The edits are based on the information shown in Supplementary Figures S2-S9. For the edits for deletions 7 and 8 no Illumina read was available. Due to the known issues of PacBio to resolve long homopolymers, we assume that the two missing nucleotides, respectively, extend the oligo-dG stretches. Low complexity regions detected by SEG during the BlastN search are shown in lower case letters. "Orig." represents the original sequence in the reference genome sequences.

#### Supplementary Figure S18 – Gene structure of *U. pustulata* (mycobiont) cox2/cob

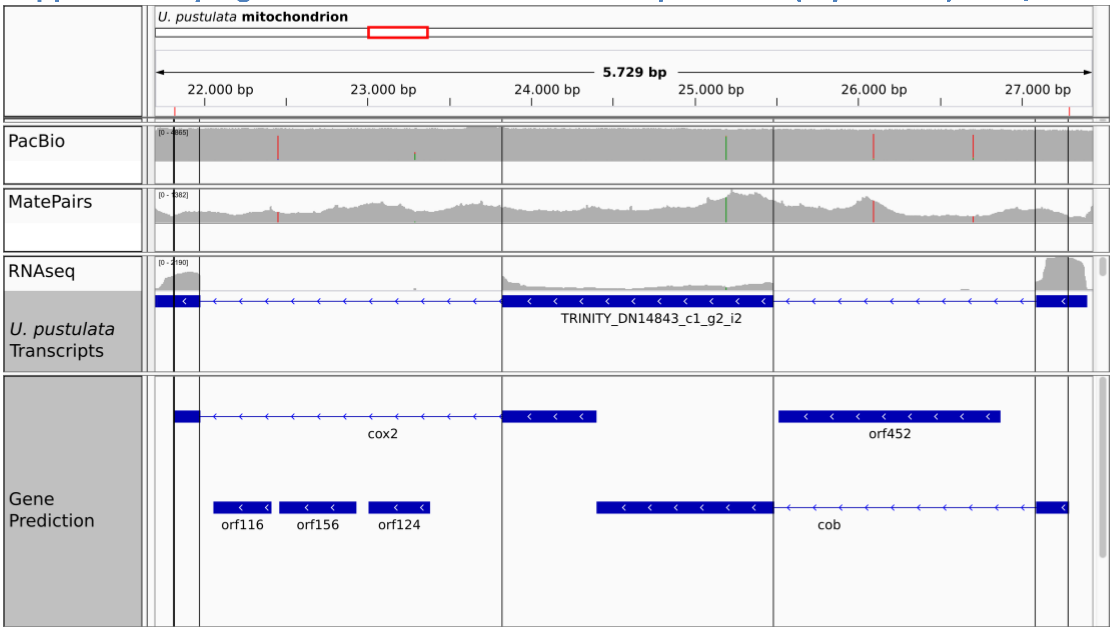

**Supplementary Figure S18 – Gene structure of *U. pustulata* (mycobiont) cox2/cob.** The gene annotation revealed that cox2 and cob reside directly next to each other in the mitochondrial genome in the same reading frame. A single transcript covers both genes, indicating that they form a single transcription unit.

#### Supplementary Figure S19 – Gene structure of *U. pustulata* (mycobiont) *nad6*

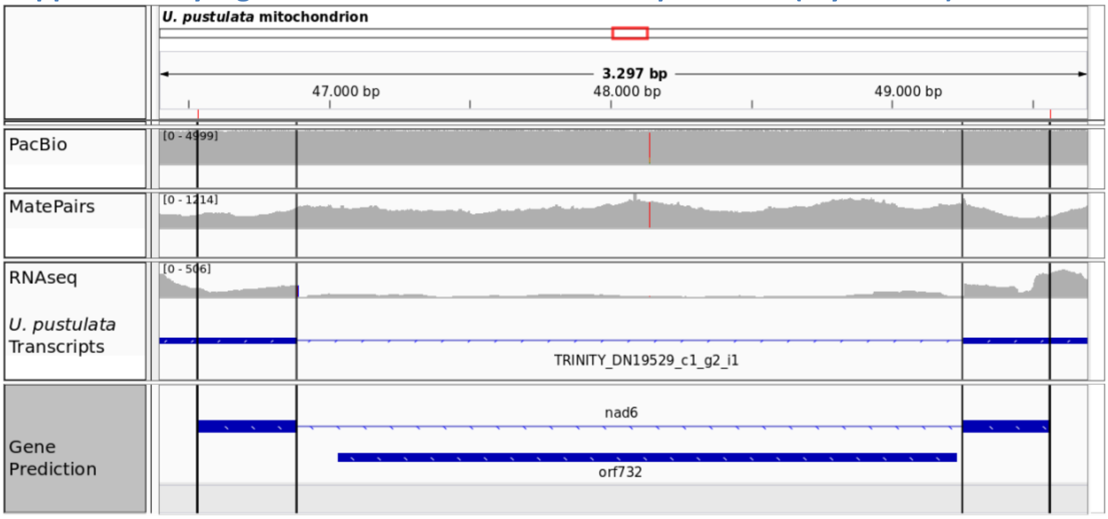

**Supplementary Figure 19 – Gene structure of *U. pustulata* (mycobiont) *nad6*.** The corresponding ORF732 encodes a 732 long putative protein harboring two Pfam domains, a reverse transcriptase domain (PF PF00078.27) and a type II intron maturase domain (PF1348.21) PFAM domain.

#### Supplementary Figure S20 – nad3 locus of *U. pustulata* (mycobiont)

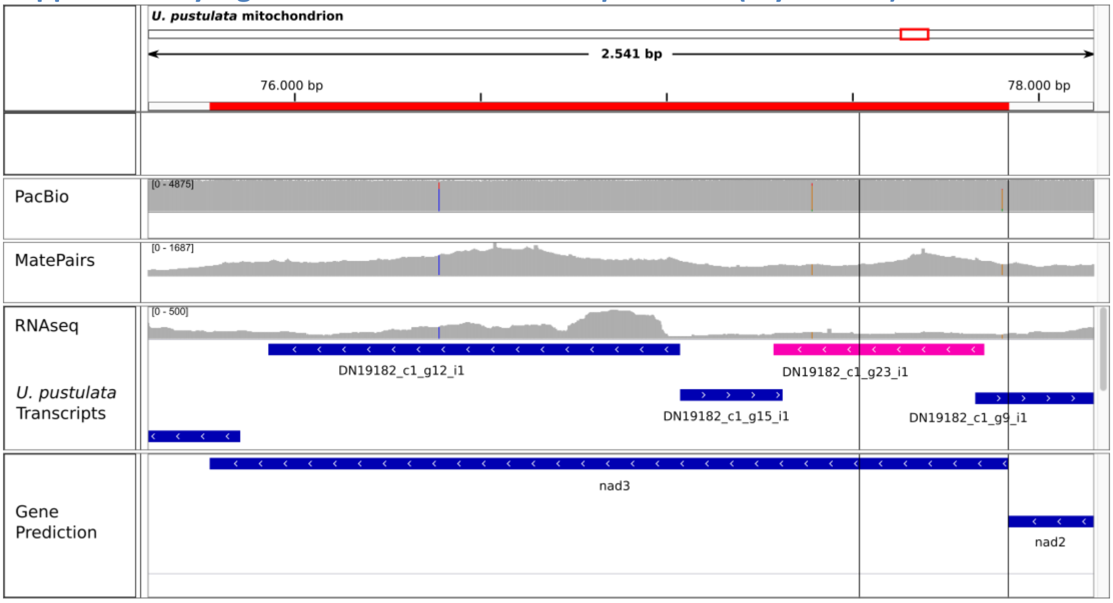

**Supplementary Figure S20 – nad3 locus of *U. pustulata* (mycobiont).** The red bar indicates the position of the nad3 ORF predicted by mfannot (track 'Annotation'). The vertical black lines delineate the begin and the end of the nad3 encoding part. The read coverage distributions of the PacBio-, MatePair-, and RNAseq libraries are provided in the corresponding tracks. The Trinity track displays transcripts that originate from the nad3 genomic region. The transcript colored in magenta covers the majority of the nad3 CDS.

#### Supplementary Figure S21 – Relocation of the chloroplast *rps4* gene to the nuclear genome of *Trebouxia sp.*

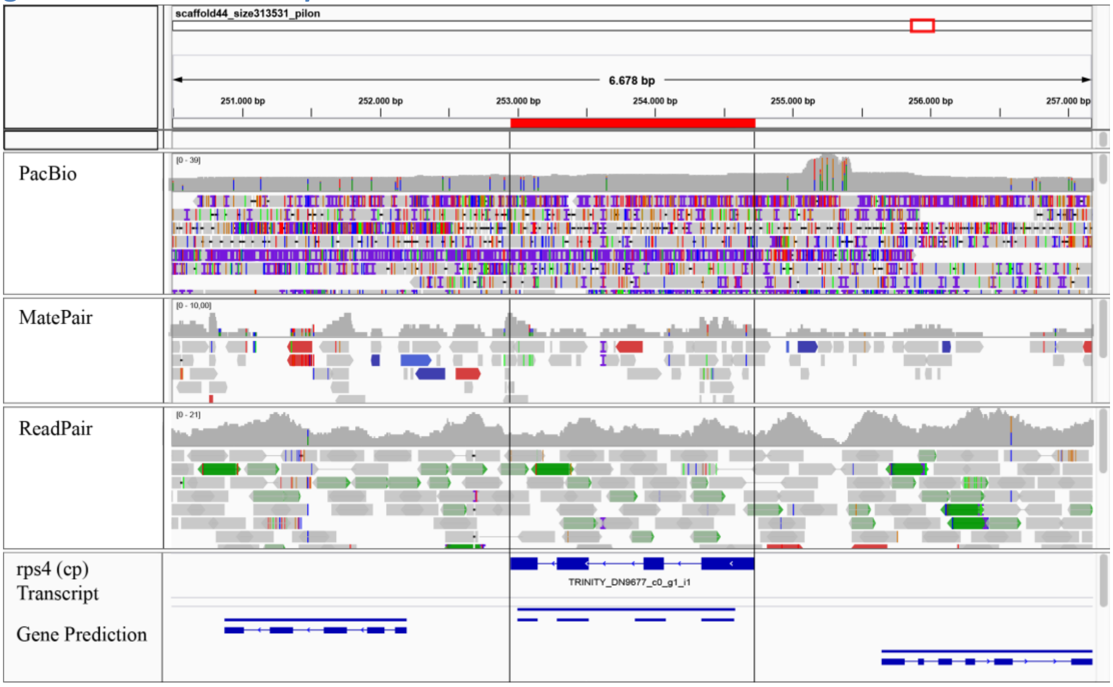

**Supplementary Figure S21 – Relocation of the chloroplast *rps4* gene to the nuclear genome of Trebouxia sp.**The rps4 locus on scaffold 44 is highlighted in red. PacBio, MatePair, and ReadPair tracks provide the read coverage distribution in this genomic region, together with the mapping positions of the individual reads. BlastX searches against the NCBI nr-prot database with the two flanking genes of rps4 reveals two nuclear genes as best hits. The gene upstream of rps4 is significantly similar to the thioredoxin, conserved site gene encoded on chromosome 10 of *Ostreococcus tauri* (XP_022839774.1). The gene downstream finds the gluthatione reductase located on chromosome 6 of *Chlamydomonas reinhardtii* as its best hit.
